# Phenotypes and Cellular Mechanics of Primary Human Aorta- and Pluripotent Stem Cell-derived Vascular Smooth Muscle Cells

**DOI:** 10.64898/2025.12.18.694995

**Authors:** Chen Yu Huang, Stanley Chun Ming Wu, Melissa Kissling, Alexander Arking, Yi-Wen Chen, Franklyn D. Hall, Shankar Sivarajan, Harrison Yezzi, Daniel H. Reich, Kenneth R. Boheler

## Abstract

**Aims:** Mature, contractile vascular smooth muscle cells (vSMC) in muscular blood vessels can de-differentiate into synthetic vSMC, while human pluripotent stem cells (hPSC) differentiate into proliferating, fetal-like vSMC that can be induced to form contractile vSMC. Given the complicated embryonic origins and conflicting views on vSMC heterogeneity, our aim was to establish how closely hPSC-vSMC of a known germ layer lineage and defined phenotype are representative of vSMC obtained from aorta (AoSMC).

**Methods and Results:** We generated paraxial mesoderm (PM)-derived synthetic and contractile vSMC from hPSC lines and performed phenotypic and mechanical comparisons with AoSMC. Synthetic hPSC-vSMC and cultured AoSMC are proliferative, sensitive to contact inhibition and exhibit phenotype switching. Contractile hPSC-vSMC are non-proliferative, capable of phenotype switching and have elevated levels of contractile proteins (CNN1, TAGLN, ACTA2). These contractile proteins also displayed density-dependent, cell-to-cell heterogeneity in both AoSMC and hPSC-vSMC. Single-cell traction force measurements on micropost array detectors (mPADs) yielded comparable forces per micropost in the nN range across cell types. Quantitative 3D force measurements on humanized smooth muscle cell microtissues (SMTs) developed similar forces per cell. Larger forces were generated by MEKi-treated hPSC-vSMC and AoSMC versus controls.

**Conclusions:** We have systematically studied the properties of hPSC-vSMC and performed comprehensive phenotypic and mechanical comparisons with AoSMC. These results demonstrate that synthetic hPSC-vSMC generated in vitro and cultured AoSMC derived from tissues are comparable and mechanically similar, but results with hPSC-vSMC are more reproducible.

**Translational Perspective:** Confirmation of how closely hPSC-vSMC of a known germ layer lineage and phenotype are representative of tissue-derived AoSMC is an unmet gap in knowledge and a requirement for their use to study human vascular (patho)physiology. We have employed paraxial mesoderm-derived hPSC-vSMC and aorta-derived vSMC (AoSMC) to compare phenotypes and to define the mechanical properties of synthetic and contractile vSMC. The results from individual vSMC and multicellular vSMC constructs with defined phenotypes confirm the adaptability of this in vitro model to reliably contribute to the investigation of human vascular pathologies.

## 1. Introduction

Vascular smooth muscle cells (vSMC) control vascular tone, vessel diameter, and blood pressure.^1^ Under physiological conditions, mature contractile vSMC are proliferatively quiescent and highly contractile. These cells express smooth muscle-prevalent proteins involved in contraction (e.g., smooth muscle actin, ACTA2; calponin 1, CNN1; transgelin, TAGLN; smooth muscle myosin heavy chain, MYH11; smoothelin, SMTN) and mediate tensile strength through interactions with the extracellular matrix (ECM). By comparison, immature synthetic vSMC in adult aorta are induced by the de-differentiation of mature, contractile vSMC or by expansion of residual populations of these cells or progenitor cells present in the tunica intima and adventitia.^2^ Synthetic vSMC are proliferative, contribute to ECM protein deposition, have relatively low levels of contractile proteins, and play a key role in the repair of injured blood vessels. Importantly, vSMC originate from multiple lineages (e.g., neural crest, proepicardium, lateral plate mesoderm, splanchnic mesoderm, paraxial mesoderm)^3, 4^ and respond in a lineage-specific way to stimuli, even when studied under identical conditions.^1, 3, 5^

The lineage origins and different functions of vSMC are key contributors to the development and progression of vascular lesions, including aortic aneurysms.^6, 7^ The heterogeneity of vSMC of different embryonic origins have a high degree of plasticity and functional differences, and the embryonic origin can affect aneurysm etiology, which differs between ascending and descending/abdominal aorta.^6, 8, 9^ Previous reports from rodent models further indicate that cultured arterial SMC are heterogeneous and show distinct phenotypic features when implanted into carotid artery.^10^ These findings suggest that SMC phenotypes are controlled not just by local stimuli and adaptations, but also by genetics, pathophysiological adaptations and embryonic origin. Moreover, the use of primary vSMC to study development or disease syndromes is complicated by differences among donors and isolation procedures. The study of vSMC from aorta (AoSMC) in vitro and their contribution to (patho)physiology is complicated by the cell’s phenotype, anatomical origin and lineage, isolation procedure, and the donors’ age and sex.

Human pluripotent stem cells (hPSC) represent an alternative, reproducible source of vSMC for in vitro modeling. Using directed in vitro differentiation protocols, vSMC derived from neural crest, lateral plate-, paraxial-, and splanchnic-mesoderm have been generated; however, detailed comparative studies are lacking that show hiPSC-derived vSMC of a known embryonic origin and phenotype retain properties that are conserved with or similar to *in vivo* derived cells.^5, 11-14^ These evaluations are critical because the origin and the regulation of phenotypic plasticity both in vitro and in vivo are likely critical determinants that contribute to the responses of vSMC to mechanical stress and vascular pathologies. In this study, we have addressed this gap in knowledge by providing systematic phenotypic and mechanical comparisons between normal human aorta-derived vSMC and paraxial mesoderm (PM)-derived hPSC-vSMC. Functionally, we assessed these cells using biophysical approaches involving quantitative 3D force measurements and complementary single-cell traction force measurements on micropillars. Despite AoSMCs showing a high degree of variability among batches, we show that AoSMC and hiPSC-vSMC both undergo phenotypic switching in vitro while retaining force generation properties that are conserved according to the phenotypic state.

## 2. Methods

Detailed experimental protocols are provided in the Supplementary material online.

### 2.1 Cell Culture

In vitro derived vSMC of PM were differentiated according to established protocols from human induced pluripotent stem cell (hiPSC) lines [JHU001/JHUi008-A (female),^15^ WTC11 (male, GM25256, Coriell Institute)] and from a human embryonic stem cell (ESC) line [H9 (female, WiCell)].^5,16^ Experiments were performed systematically on JHU001-vSMC, with experiments from other lines used to identify possible clonal line variations.^17^ Experiments were performed on both PM-derived vSMC enriched with lactate-medium^18^ and on primary AoSMC obtained from commercial vendors (See *Table 1*). In most experiments, synthetic hPSC-vSMC were treated with vehicle (DMSO) or 1 mM of the mitogen-activated protein kinase kinase inhibitor (MEKi) PD0325901to induce a contractile phenotype.^11^

**Table 1:** Commercially obtained batches of aorta-derived vSMCs.

| Reference Name | Company | Line/Batch Number | Gender | Age (years)/Race | Passage | Doubling Time (hours) | Donor Source (likely mixed thoracic + abdominal) |
| --- | --- | --- | --- | --- | --- | --- | --- |
| **AoSMC1** | ATCC | 70045161 | Male | 48/Caucasian | p3 | 26 | Aorta |
| **AoSMC2** | ATCC | 70008916 | Male | 34/White | p2 | unknown | Aorta |
| **AoSMC3** | Lonza | 21TL316170 | Female | 24/Hispanic | p3 | 22 | Aorta |
| **AoSMC4** | Lonza | 18TL117408 | Female | 11/Black | p3 | 38 | Aorta |
| **AoSMC<sub>f</sub>** | ScienCell | 29308 | unknown | 18-week gestation/<br>unknown | unknown | unknown | Aorta (thoracic + abdominal) |

### 2.2 Phenotypic Analyses of vSMC

RNA was isolated using the RNeasy Plus Mini Kit (Qiagen) and cDNA generated using the High-Capacity cDNA Reverse Transcription Kit (ThermoFisher). Amplifications were performed with the ViiA^TM^ 7 Real-Time PCR System. Data were normalized to RPL32.^19^ Some qPCR data are reported with TaqMan probes (Supplementary material online, *Tables S1a, S1b*). Total DNA was obtained from lysed cells^19^ and incubated with 1X PicoGreen according to the manufacturer’s instructions (ThermoFisher Scientific, USA). Fluorescence was measured at Ex485 nm/Em528 nm, and DNA content determined relative to salmon sperm DNA standards. Cell viability was assessed using PrestoBlue (ThermoFisher) and the LIVE/DEAD™ Cell Imaging Kit (ThermoFisher). Cell proliferation was evaluated from cell numbers as a function of time. Cell cycle analyses were performed from TrypLE digested cells that were fixed, incubated with propidium iodide (50 μg/mL PI, Sigma) and assessed by flow cytometry. Cell cycle compartments were deconvoluted from single-parameter DNA histograms of ∼10,000 cells.

Protein lysates were isolated in radio immunoprecipitation assay (RIPA) buffer (Millipore Sigma, USA), supplemented with phosphatase and protease inhibitors (Cell Signaling Technology, USA), and protein content quantified using the BCA protein assay (ThermoFisher). Following electrophoresis on SDS-PAGE gels and transfer, PVDF membranes were incubated with primary and secondary antibodies (Supplementary material online, *Table S2*). Images were captured (BioRad ChemiDoc Touch Imaging System), and data quantified using ImageJ. Coomassie blue stained membranes showing total protein loading or β-tubulin signals were used to normalize data. Whole gel images are available in Supplementary material online (*Other*).

For immunostaining, cells were cultured overnight or to confluency. Cells were washed with DPBS, fixed with 4% paraformaldehyde, permeabilized in 0.1% Triton in PBS, and incubated in a 1% bovine serum albumin solution. Primary antibodies were added to cells overnight, washed three times, and incubated with secondary antibodies for 1 hr, followed by addition of DAPI (1:200; ThermoFisher). Images were acquired using the Zeiss LSM 780 Confocal Laser Scanning Microscope (Carl Zeiss AG) and analyzed with the corresponding software Zen (Carl Zeiss AG). For immunophenotyping by flow cytometry, cells were fixed and permeabilized using the FIX & PERM Cell Fixation and Cell Permeabilization kit (ThermoFisher) and incubated with primary and then secondary antibodies. For antibody details, see Supplementary material online, *Table S2*.

### 2.3 Mechanics of vSMC

Using micropost array detectors (mPADs),^20^ we measured the average traction force generation of single hPSC-derived vSMC from images of deflected microposts in contact with the cells, taken ∼24 hours after plating cells onto the devices. Individual microposts in an mPAD report a force 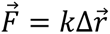 where 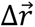 is the deflection of the micropost’s tip from its resting position, and *k* is the posts’ spring constant for small lateral deflections. In these studies, *k* = 15.7 nN/µm.^21, 22^ We quantified the force generated by each cell via the average of the magnitude of the traction forces, *F_avg_*, over the set of *N_posts_* posts to which the cell was adhered,^22^

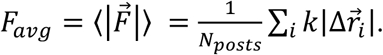

This provides a measure of cellular contractility, reported as the force per post.

We constructed smooth muscle microtissues (SMTs) in arrays of PDMS microwells with microfabricated tissue gauges (µTUGs) containing a pair of flexible pillars spaced by 500 µm.^23,24^ Cells were dissociated with TrypLE, and mixtures of vSMC or AoSMC with ECM (human-sourced collagen and fibrinogen) were seeded and allowed to compact to form a microtissue structure spanning the pillars. The pillars act like simple springs for small deflections (*k =* 0.25 µN/µm or *k =* 0.5 µN/µm) and report the collective force generated by the ∼100 cells in the SMT. Data are presented as total SMT force, as forces normalized per cell based on a count of nuclei in each tissue using DAPI, and as force per cross-sectional area (SMT stress) to correct for variability in SMT force arising from variations in SMT size.

### 2.4 Data Analysis and Statistics

Flow cytometry analysis was performed on a CytoFLEX (Beckman Coulter) and analyzed (FlowJo, LLC). For mPADs, videos were processed^25^ in Igor Pro 9 (WaveMetrics) to determine the displacement of each post beneath the cell from its resting position and to calculate *F_avg_* for each cell. Data are presented as mean ± standard deviation (SD) or mean ± standard error (SEM). Statistical tests included one-way and two-way ANOVA, and a two-sample Student’s *t*-test. Statistical differences were considered significant at p < 0.05. Significance was determined using a Welch’s (unequal variance) *t-*test if the data failed to meet the assumptions of a Student’s *t*-test.

## 3. Results

### 3.1 Differentiation and Enrichment of human Pluripotent Stem Cell derived – vSMC

Human pluripotent stem cells (hPSC) were differentiated as shown schematically in *Figure 1A*. At differentiation day (DD) 5, transcripts encoding the T-box transcription factor 6 (TBX6) increased by 18.1±0.03-fold (p<0.05, n=3) relative to undifferentiated JHU001 hiPSC, confirming the induction of paraxial mesoderm. Representative images of undifferentiated hPSC and differentiating progeny are provided (Supplementary material online, *Figure S1A)*. Brightfield imaging of sub-confluent cells showed mixed rhomboid- and fibroblast-like (spindle-shaped with cellular processes) morphologies (Supplementary material online, *Figure S1A*, DD18).^26^ When confluent, the cells developed a spindle-shaped (fusiform) morphology (*Figure 1B*, SMCM and Supplementary material online, *Figure S1B*). These cell density-dependent morphologies were similar across all tested hPSC lines at passages 1-7.

**Figure 1.**
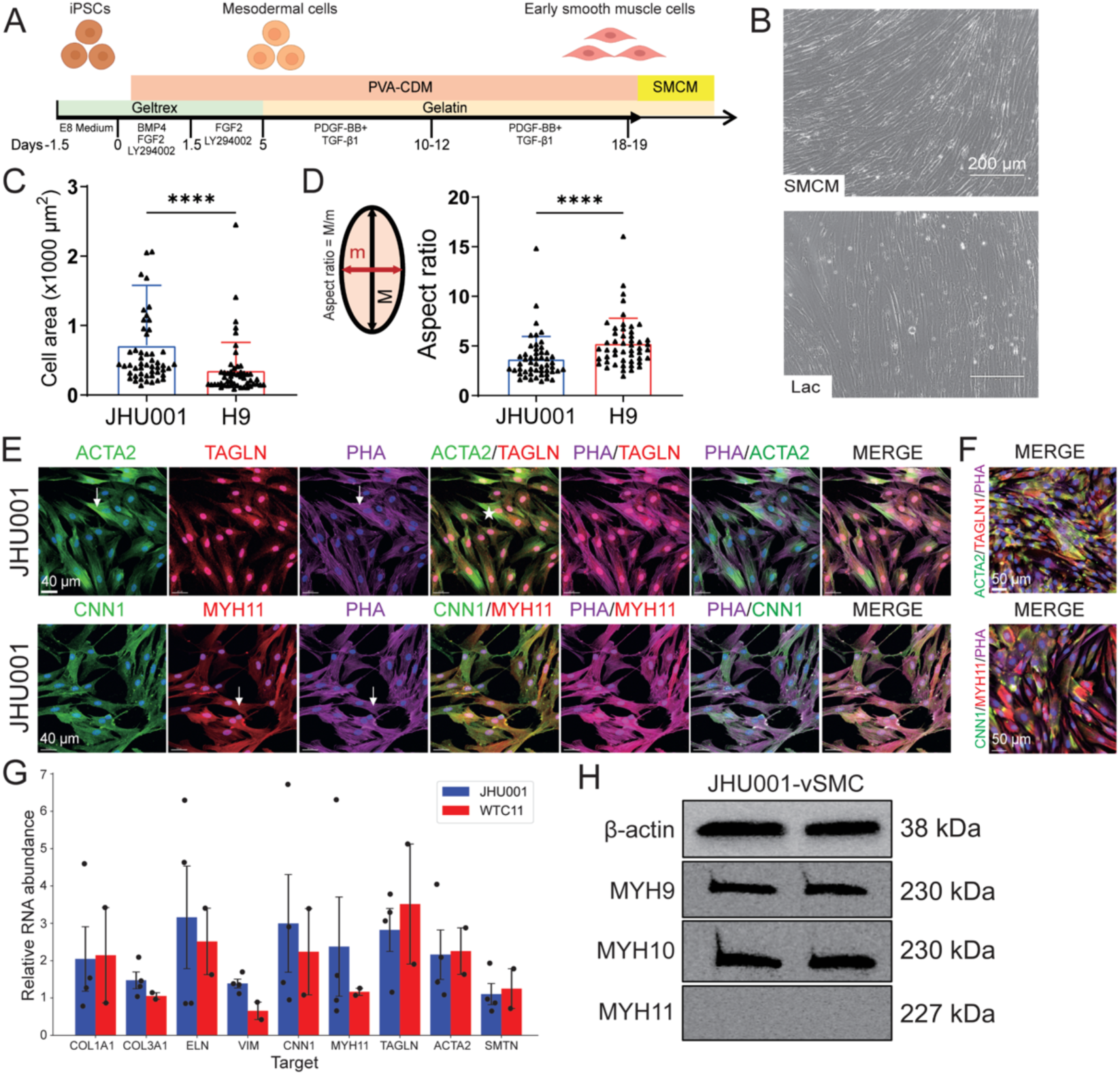
Characteristics of hiPSC-vSMC. A) Schematic showing the generation of PM-derived vSMC from hPSC. B) Brightfield images of confluent JHU001-vSMC cultured in SMCM and in Lac medium for 2 days. Scale bar 200 μm. Physical characteristics of PM-derived hPSC-vSMC from two lines are shown in C) cell area and D) elongation (aspect ratio of best-fit ellipse). Error bars: ±SD. JHU001: n=51; H9: n=50. E) Immunostaining of sub-confluent cultures of JHU001-vSMCs. White stars indicate vSMC with differential ACTA2 and TAGLN signal intensity. White arrows show filamentous-like structures. Scale bar 40 μm. F) Immunofluorescent (IF) images of confluent cultures of JHU001-vSMC showing heterogeneous staining in merged images. Scale bar 50 μm. G) qPCR data of transcripts from hPSC-derived vSMC normalized to RPL32. Error bars: ±SEM, n=3-4. H) Composite figure showing bands from Western blots for β-actin, MYH9, MYH10, and MYH11 (1s exposures) from lysates prepared from two independent batches of JHU001-vSMC. Statistics were performed using a Student’s *t*-test in E and F, a Welch’s *t*-test in G, and a two-way ANOVA in H. *P*<0.05 was considered statistically significant (*p<0.05, **p<0.01, ***p<0.001, ****p<0.0001).

Glucose-free, lactate-containing medium has been reported to enrich populations of muscle cells and to select for more synthetic vSMC.^27^ We incubated near-confluent PM-derived hPSC-vSMC with SMCM or RPMI medium supplemented with either 4 mM lactate, hFGF2 and hEGF (Lac + GF) for 6 days as described^27^ or DMEM supplemented with 4 mM lactate and lacking any additional growth factors (Lac) for 2 to 4 days.^18^ In brief, cultivation of the cells with Lac + GF for 6 days often led to over-confluent cultures, while Lac treatments for 4 days led to extensive cell death. Treatment of cultures with Lac for 2 days resulted in confluent cultures of fusiform-shaped vSMC with a minimal number of contaminating non-vSMC (*Figure 1B,* Supplementary material online *Figure S1B*). Comprehensive comparisons are provided (see Supplementary material online, *Results and Figures S1B*-*S5)*. Overall, the enrichment process did not lead to any substantive phenotypic changes relative to non-enriched controls. All subsequent analyses were performed on hPSC-derived vSMC selected for 2 days with Lac medium and re-fed with SMCM for 1-2 days prior to experimentation.

### 3.2 PM-vSMC synthetic phenotype

We systematically characterized the Lac-enriched PM-vSMC to establish phenotypes and to address heterogeneity. At low densities, we observed mixed rhomboid and fibroblast-like vSMC; but in confluent cultures, the cell shape was fusiform. Morphometrically, the cell areas of JHU001-vSMC were larger than H9-vSMC (*Figure 1C*). The aspect ratio of both JHU001- and H9-vSMCs were consistent with an elongated vSMC morphology, although that of the H9-derived vSMCs was measurably larger (*Figure 1D*). By immunostaining (*Figure 1E*), we observed relatively uniform signals for TAGLN, MYH11, CNN1 and F-actin (phalloidin, PHA) in hPSC-vSMC. However, signal intensities for cells co-stained with ACTA2 and PHA antibodies varied (*Figure 1E*), and signals for contractile ACTA2 were stronger in some TAGLN-positive cells (See star symbols, *Figure 1E*). Filamentous-like structures were observed in cells immunostained for PHA, ACTA2 and MYH11 (see white arrows, *Figure 1E*), while TAGLN fluorescence was generally diffuse.

At high densities, vSMC marker immunostaining was heterogeneous when compared to low-density cultures (*Figure 1F*, Supplementary materials online, *Figures S2A and S2B*). The fluorescence signals for all proteins analyzed were variable, indicating that cell densities and time of growth may affect cell phenotypes. However, RNA from JHU001- and WTC11-vSMC cultures did not show significant differences for any transcript encoding extracellular matrix, intermediate filament, actin-interacting, contractile or structural protein (*Figure 1G*). Similar results were observed with H9-vSMC for transcripts quantified with Taqman probes (not shown). By Western, ECM and vSMC proteins were present in similar amounts in JHU001 and WTC11 vSMC (Supplementary material online, *Figure S4 and S6D*). We observed by flow cytometry that the majority of JHU001-vSMC were positive (90-95%) for CNN1, TAGLN and ACTA2 when compared to antibody isotype controls or unstained cells (Supplementary materials online, *Figures S5A-S5D)*. The relatively broad histogram peaks for these proteins indicate a degree of heterogeneity (signal variation) within these cell populations. In WTC11-derived vSMC, the number of CNN1^+^, ACTA2^+^ and TAGLN^+^ cells ranged from 90-98% relative to isotype controls (Supplementary materials online, *Figures S5E, S5F)*.

We observed strong fluorescent signals for MYH11, a marker for the contractile phenotype, in fixed cells cultivated in SMCM (immunostaining, *Figure S2A*; flow cytometry (not shown)), which raised questions regarding antibody specificity. We therefore quantified transcripts encoding non-muscle myosin heavy chains and MYH11 by qPCR. Transcripts encoding MYH9 (Ct_avg_=17.1±0.4) and MYH10 (Ct_avg_=18.8±0.7), but not MYH11 (Ct_avg_=28.6±2.4) were abundant in JHU001-vSMC (Supplementary material online, *Figure S6A*). Similarly, by Western blotting, MYH9 and MYH10 proteins were detected after short exposures (∼1 sec), while MYH11 was detectable after longer exposures (∼60 sec) (*Figure 1H*, Supplementary material online, *Figure S6B*). Similar results were observed in H9-vSMC (Supplementary material online, *Figure S6C*). These Westerns are consistent with either an absence or low levels of MYH11 in hiPSC-vSMC cultivated in SMCM. Consequently, the fluorescent signals observed with this antibody in fixed cells likely reflect cross-reactivity with other myosin proteins (MYH) or non-specific binding.

### 3.3 PM-vSMC contractile phenotype

Kumar et al. reported that MEKi could induce mesenchymoangioblast-derived synthetic vSMC (i.e., splanchnic mesoderm) to form a more mature, proliferatively quiescent, contractile phenotype.^11^ We show that enriched PM-vSMC could also form a contractile phenotype after treatment with PD0325901 (MEKi). Inhibition led to distinctly different growth characteristics of hPSC-vSMC relative to DMSO controls. Structurally, the MEKi-treated cells had lengths and widths larger than DMSO-treated controls. The overall cell density decreased after 6 days of treatment (*Figure 2A*, Supplementary materials online, *Figure S7*). When MEKi-treated cells were dissociated and plated as single cells, low-density vSMC assumed rhomboid or fusiform shapes (*Figure 2B* for JHU001 and Supplementary materials online *Figure S8A* for H9), with cell sizes in two dimensions 2- to 3.5-fold greater than controls for both JHU001- and H9-vSMC. Control vSMC generally had lower signal strengths for CNN1 than MEKi-treated cells. The latter developed internal fiber-like structures associated with a more mature phenotype (*Figure 2B*). The cell area increased significantly, but we could not demonstrate any differences in the nuclear size for JHU001-vSMC (*Figure 2C*). However, MEKi increased the nuclear area relative to controls (p<0.0001) for H9-vSMC. The aspect ratios (*Figure 2C*) also significantly decreased with MEKi treatment, consistent with the formation of a more symmetric (less spindle-shaped) cell type. (*Figure 2C, and* Supplementary materials online *Figure S8B*).

**Figure 2.**
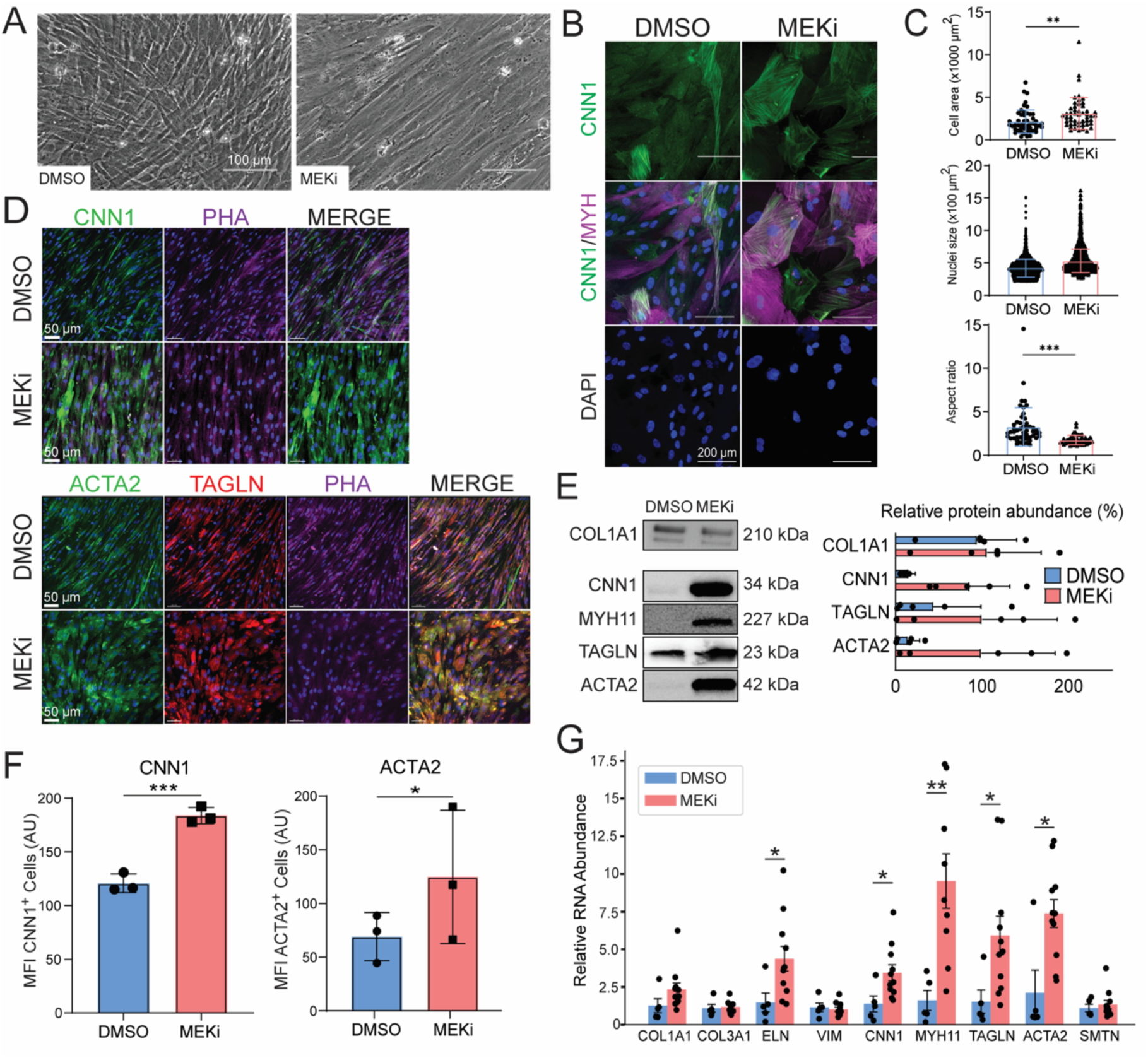
JHU001-derived vSMC treated with DMSO or MEKi. A) Brightfield image of monolayer cultures following treatments. Scale bar 100 μm. B) IF images of replated cells after 6-day treatments, 1-2 days of recovery, and passaging onto coverslips. Cells were fixed and stained as indicated (MYH IF was considered non-specific). Scale bar 200 μm. C) Physical characteristics of control and MEKi-treated cells. Aspect ratio is defined as in Figure 1D. Error bars: ±SD, DMSO n=50, MEKi n=50. D) IF images of vSMC contractile protein markers including CNN1, ACTA2, TAGLN and F-actin in monolayer cultures. Scale bar 50 μm. E) Western blots of selected proteins in hPSC-vSMC ± MEKi. Graphed data from one gel (COL1A1) and another gel (CNN1, MYH11, TAGLN, ACTA2) were normalized to total protein loading and plotted as a percentage; however, exposure times varied. Error bars: ±SD, n=5. F) Mean fluorescence protein intensity (MFI) of selected proteins in MEKi-versus DMSO-treated cells evaluated by flow cytometry. G) qPCR analysis for the indicated transcripts. Error bars: ±SEM, n=5-11. Error bars: ±SD, n=3. Statistics were performed using a Student’s *t-*test in C, a Welch’s *t-*test in D, a two-way ANOVA in E and a one-way ANOVA in G. *P*<0.05 was considered statistically significant (*p<0.05, **p<0.01, ***p<0.001, ****p<0.0001).

In high-density MEKi-treated cultures, the fluorescent signal intensities for CNN1, ACTA2 and TAGLN were stronger when compared with vehicle-control cells. Filamentous-like protein structures also were more readily observed (*Figure 2D*, Supplementary materials online, *Figure S8C*). A significant increase in contractile protein content was confirmed by Western blotting (*Figure 2E*). When ACTA2 and CNN1 were examined by flow cytometry, significantly higher mean fluorescence protein intensities (MFI) were observed in MEKi-treated relative to vehicle-control cells (*Figure 2F*). At the RNA level, MEKi treatment of confluent cells led to significant increases in ELN, CNN1, MYH11, TAGLN and ACTA2 transcripts in JHU001-vSMC relative to controls (*Figure 2G*). As a final comparison, experiments were performed with rapamycin, which has been reported to cause a contractile phenotype in human and rodent vSMC.^28^ In our experiments, this treatment resulted in cultures with spindle-shaped cells similar to controls (Supplementary materials online, *Figure S7*), but it did not induce the expression of contractile proteins or affect cell mechanics (not shown).

### 3.4 Cell proliferation and cell cycle analyses

Our data are consistent with the in vitro differentiation of hPSC-vSMC having a synthetic-like phenotype; however, MEKi-treated contractile-like cells were visually larger than the DMSO-treated vSMC, possibly due to changes in cell density, proliferation or cell viability. Under control conditions, hPSC-vSMC initially doubled approximately every 2-3 days, with a ∼4-5-fold increase over 10 days (*Figure 3A*). As the cells reached confluency, the rate of change decreased. Consistently, the DNA content of control hPSC-vSMC increased as a function of time (*Figure 3B*). Anecdotally, if the original plating density was semi-confluent or confluent, the vSMC number would only increase by ∼3- or 1.5-fold, respectively over the course of one week (not shown). By comparison, the total number of MEKi-treated cells plated at a sub-confluent density increased by ∼1.5-fold between day 0 and day 2 of cultivation, after which time the cell numbers plateaued or decreased (*Figure 3A*, n=6, p<0.05 for MEKi versus Ctl). The DNA content did not differ relative to controls after 2 days with MEKi, but it decreased between days 4 and 8 (p<0.05) (*Figure 3B*). By PrestoBlue staining, a transient decrease in overall viability of MEKi-treated vSMC occurred between days 2 and 5. No additional change in viability was observed at later times (*Figure 3C*), and no difference in live versus dead JHU001-vSMC numbers could be demonstrated after 6 days of DMSO or MEKi treatment (*Figure 3D, p>0.05*).

**Figure 3.**
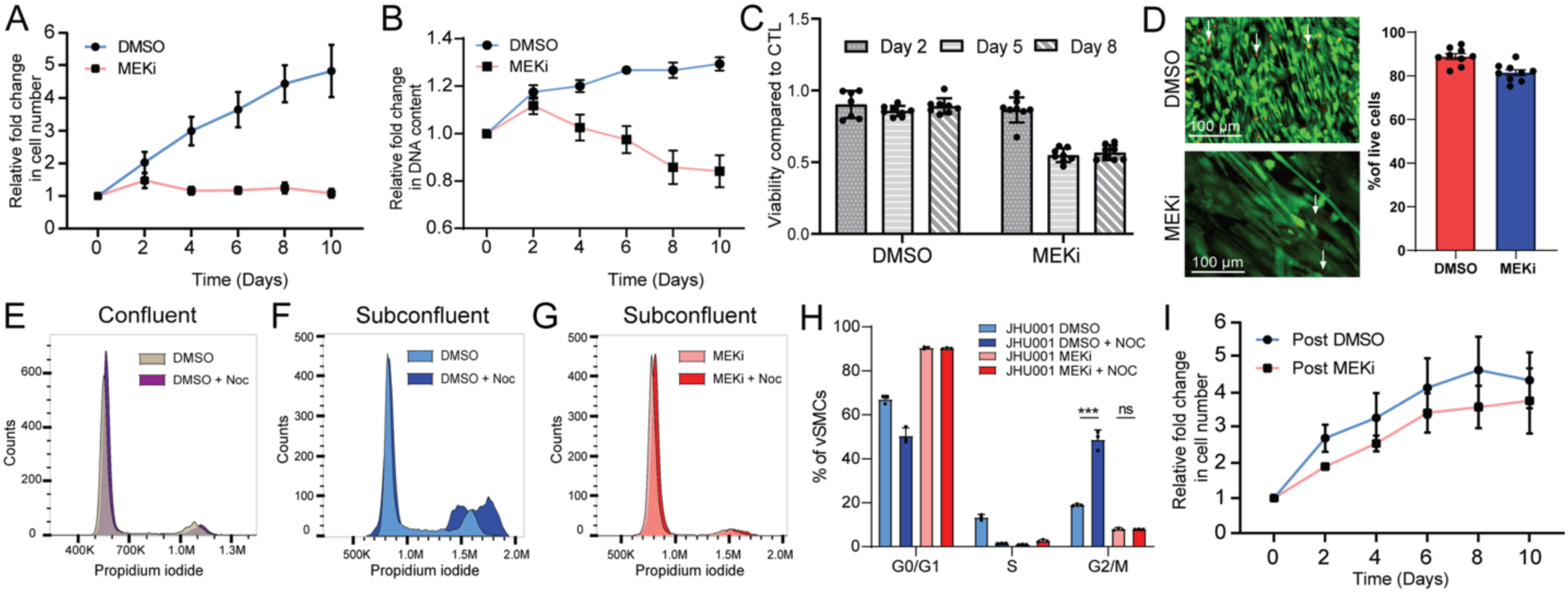
JHU001 hPSC-vSMC proliferation, viability, and cell cycle attributes. A) Cell counts as a function of time showing fold changes during cultivation with MEKi versus control (Error bars: ±SEM, n=6, p<0.005 for D4 onward). B). DNA content measured by PicoGreen assays with time. (Error bars ±SEM n=6, p<0.005) C) Cell viability assays performed with PrestoBlue, showing that the viability of MEKi-treated cells with time. (Error bars: ±SD, n=7-8). D) Live-Dead cell staining (as a percent) after 6 days of MEKi treatment and 2 days of recovery. Dead cells are indicated by white arrows. Scale bar 100 μm. Representative histograms showing cell cycle phases of E) confluent hPSC-vSMC cultures ± Noc and F) subconfluent cultures of vSMCs ± Noc. G) Cell cycle phases of subconfluent cultures of MEKi-treated vSMC ± Noc. H) Graphic representation quantifying the percentage of subconfluent vSMC in each phase of the cell cycle with treatments as indicated. Error bars: ±SD, n=3. I) Proliferation of contractile vSMC following withdrawal of MEKi and replating of cells. Error bars: ±SEM, n=6. Statistics were carried out using a Student’s *t-*test in A, a two-way ANOVA in B, and a one-way ANOVA in C and H. *P*<0.05 was considered statistically significant (**p<0.01, ***p<0.001, ****p<0.0001).

We performed cell cycle analyses on fixed cells incubated with PI, a double-stranded DNA intercalating dye (*Figures 3E-3G)*. Under confluent control conditions, the proportion of vSMC (JHU001) in the G0/G1 phase was 81±3%. Only 11±2 % of the cells were in the S phase, with the remaining 10±2% of vSMC in the G2/M phase. After 24 hours with nocodazole (Noc), a microtubule inhibitor that causes cell cycle arrest mostly in G2/M, the number of cells in G0/G1, S and G2/M did not significantly change (G0/G1: 79±4%; S: 6±1%; G2/M: 14±1%) relative to controls. These data are consistent with contact inhibition of proliferation of vSMC. Therefore, we passaged hPSC-vSMC and repeated the experiments on sub-confluent cells. Under these conditions, 67±2% of control JHU001-vSMC were in the G0/G1 phase, 13±1% in the S phase, and 19±0.4% in the G2/M phase. Following 48 hours of Noc treatment, 50±4% of the vSMC were still in G0/G1, and 49±4% were in G2/M (*Figures 3F, 3H).* We observed two peaks in the G2/M phase, indicative of aneuploidy likely caused by mitotic slippage following Noc treatments. Similar results were observed with WTC11-vSMC (Supplementary materials online, *Figure S9*).

Next, we analyzed the cell cycle of MEKi-treated cells plated at sub-confluent plating densities. When JHU001-vSMC were cultured for 48 hours, 90±0.6% of the vSMC were in G0/G1, and only 1±0.1% and 8±1% of the cells were in the S and G2/M phases of the cell cycle respectively (*Figures 3G, 3H*). Following Noc treatment, the proportion of cells in these three phases of the cell cycle did not significantly change, and little to no aneuploidy was observed. Data for WTC11-derived vSMC were similar (Supplementary materials online, *Figure S9*). Thus, MEKi-treated cells had either withdrawn from the cell cycle (G0) or experienced a G1 checkpoint block. However, when these cells were passaged at sub-confluent densities and cultured in SMCM for 10 days, the vSMC proliferated to become confluent and could be passaged (*Figure 3I*).

### 3.5 Aorta-derived vSMC

To compare hPSC-derived vSMC with patient-derived vSMC, we purchased four batches of human adult aorta vSMC (AoSMC), and one batch of fetal aorta (AoSMC_f_)(*Table 1*). The donor information and doubling times for three of the batches were provided by the suppliers. In our hands, AoSMC cultured in SMCM proliferated and could be passaged; however, the time needed to reach confluency increased from ∼7-10 days to >10-18 days as the passage number increased.

Morphologically, AoSMC are heterogeneous. AoSMC1 and AoSMC2 cultures contained rhomboid, polygonal, and fibroblast-like cells. AoSMC3 cultures had mostly rhomboid cells and fibroblast-like cells, while AoSMC4 exhibited diverse morphologies (*Figure 4A*). AoSMC_f_ cultures consisted mostly of spindle-shaped cells at high densities and rhomboid or fibroblast-like morphologies at low densities (*Figure 4A*). Two days of cultivation of AoSMC in Lac medium did not substantially change cell numbers or morphologies except in AoSMC4 (not shown).

**Figure 4.**
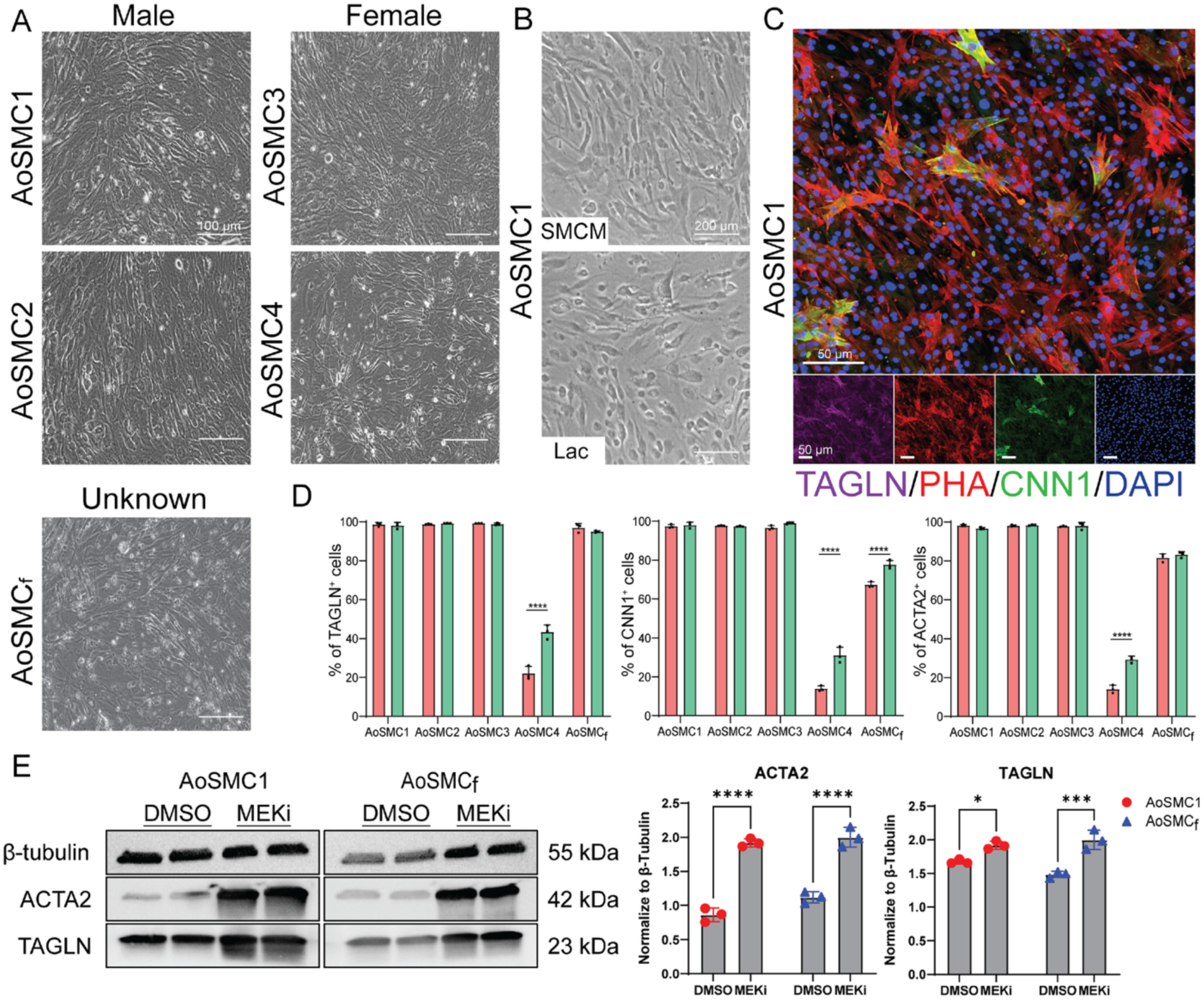
AoSMC batch comparisons. A) Brightfield images of AoSMC and AoSMC_f_ lines. B) Brightfield images of AoSMC1 incubated for 2 days with Lac vs SMCM. C) CNN1, TAGLN and F-actin (phalloidin (PHA)) immunostaining and DAPI staining of confluent AoSMC. The individual color channels are shown below the image. Scale bar 50 μm. D) Flow cytometry analysis showing the percentage of cells positive for the indicated marker proteins. Error bars: ±SD, n=3. E) Representative Western blots of indicated proteins in AoSMC ± MEKi. Data obtained from independent lysates were normalized to β-tubulin and graphed. Error bars: ±SD, n=3. Statistics were carried out using a two-way ANOVA in D and E. *P*<0.05 was considered statistically significant (*p<0.05, ***p<0.001,****p<0.0001)

By immunostaining, the fluorescent signal intensity and cell-to-cell signal heterogeneity varied across the AoSMC lines (Figure 4C, Supplementary materials online, *Figure S10*). CNN1 in AoSMC2 and AoSMC3 showed cell-to-cell variability, with generally diffuse signals. ACTA2 and TAGLN signal intensities were stronger, with heterogeneous filamentous-like structures observed in AoSMC1 and AoSMC3 when compared to AoSMC2. ACTA2 and TAGLN formed fiber-like structures that were prominent in larger cells present in AoSMC1 and AoSMC3 cultures. AoSMC_f_ generally had weak ACTA2 and CNN signals, with some filamentous-like structures in a subset of these cells.

We evaluated the AoSMC lines and AoSMC_f_ ± Lac medium by flow cytometry (*Figure 4D*) and qPCR. The number of cells positive for CNN1, ACTA2 and TAGLN ± Lac medium ranged from ∼90-98% and did not differ among AoSMC1, AoSMC2, and AoSMC3. However, a fourth batch, AoSMC4, was only ∼10%, ∼10% and ∼20% positive for CNN1, ACTA2, and TAGLN, respectively (*Figure 4D*). After cultivation in Lac medium, TAGLN, ACTA2 and CNN1 positivity increased to ∼40%, ∼30%, and ∼30%, respectively. Due to the relatively high proportion of vSMC marker-negative cells (i.e., non-vSMC), no additional experiments were performed on AoSMC4. In AoSMC_f_, TAGLN was present in ∼95% of the cells, while CNN1 and ACTA2 were observed in ∼65%, and ∼80% of the cells, respectively. Treatment with Lac medium increased the percentage of CNN1^+^ cells to ∼75% and significantly increased transcripts encoding CNN1, TAGLN and ACTA2 relative to controls (*Figure 4D*, Supplementary materials online, *Figure S11*). AoSMC1-3 also showed significant increases in CNN1 and TAGLN RNA as well as other selective changes in transcripts relative to controls. In summary, Lac medium did not enrich for synthetic AoSMC. Instead, it often led to increases in transcripts encoding contractile proteins. For this reason, no additional enrichments with Lac medium were performed on any of the AoSMC.

As a final test, we determined whether MEKi could induce a more contractile-like phenotype in AoSMC. By Western blotting, ACTA2 and TAGLN proteins were significantly increased following MEKi treatment in both AoSMC1 and AoSMC_f_ (*Figures 4E, 4F*, n=3); however, the increase in TAGLN was much more significant in AoSMC_f_. Force measurements of MEKi-treated AoSMC also generally increased relative to controls (see below), with the exception of AoSMC_f_, confirming that MEKi can induce a more contractile phenotype in these cells.

### 3.6 Force measurements of hPSC-vSMC and AoSMC

Cell force measurements using microfabricated post-array detectors (mPADs) and humanized vSMC microtissues were employed to quantify traction forces exerted by cells on a substrate and to study collective cell forces in a tissue model, respectively. Experiments were performed ± MEKi with enriched hPSC-vSMC and AoSMC positive for vSMC markers.

#### 3.6.1 mPADs

Using mPADs, we measured the average traction force generation of single hPSC-derived vSMC from images of deflected microposts in contact with the cells as shown schematically in *Figure 5A*. Examples of traction force vector maps for a JHU001-vSMC and an AoSMC are shown in *Figure 5B*. The average traction force *F_avg_* for each cell typically fell in the range 1-4 nN/post (*Figure 5C*). Proliferative WTC11-vSMC showed significantly larger forces than JHU001-vSMC, and both lines showed significant increases in *F_avg_* with MEKi treatment relative to DMSO controls. Variations in *F_avg_* between independent differentiation batches (n=4, Supplementary materials online, *Figure S12*) were somewhat larger for the WTC11-vSMC, but these results show that MEKi treatment reliably yields greater contractility, consistent with the increase in markers for the contractile phenotype reported above.

**Figure 5.**
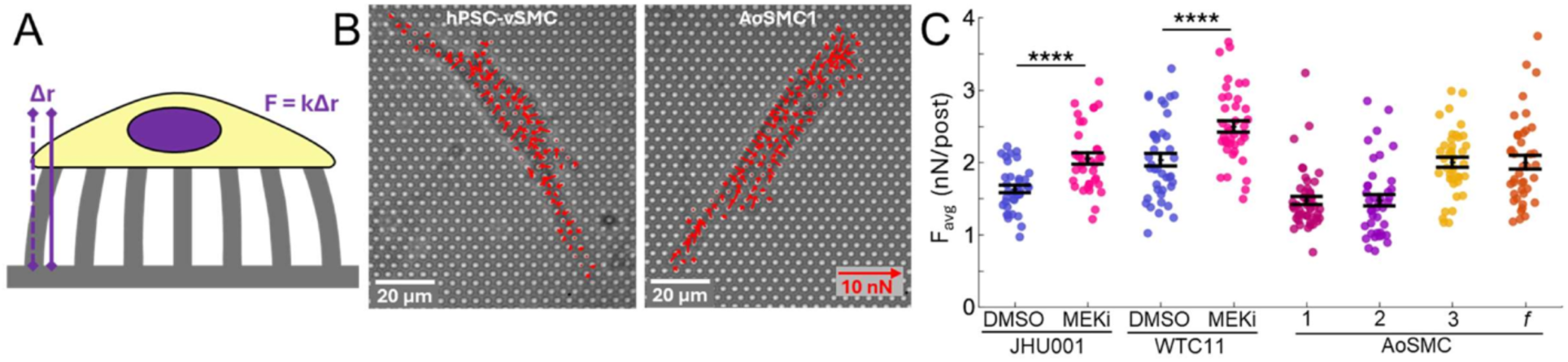
Measurement of single vSMC baseline contractile properties using mPADs. (A) Schematic of a cell on a mPAD substrate, showing deflection of the flexible microposts used to measure the cell’s force generation. (B) Representative images of an hPSC-derived vSMC and an AoSMC on mPADs showing vector forces (arrows). The posts were 1.8 µm in diameter and spaced on a 4 µm pitch. Scale bar 20 μm. (C) Average force per post *F_avg_* for (i) JHU001-vSMC: 1.63±0.05 nN/post (DMSO, n=36 cells), 2.05±0.08 nN/post (MEKi, n=40 cells); (ii) WTC11-vSMC 2.04±0.09 nN/post (DMSO, n=40 cells), 2.04±0.09 nN/post (MEKi, n=40 cells)); and (iii) AoSMC1: 1.47±0.06 nN/post, (n=50 cells); AoSMC2: 1.47±0.08 nN/post, (n=40 cells), AoSMC3: 2.00±0.07 nN/post, (n=40 cells, and AoSMC_f_: 2.0±0.1 nN/post (n=40 cells). Error bars: ±SEM. Statistical analysis was carried out using a Welch’s *t-*test in C for the DMSO versus MEKi groups. *P*<0.05 was considered statistically significant (****p<0.0001).

Measurements of AoSMC force using mPADs showed greater variations than hPSC-vSMC (*Figure 5C*). Variations were observed among distinct batches of cells and with passage number (Supplementary materials online, *Figure S12C*). Overall, a pooled comparison of the results in *Figure 5C* show that MEKi-treated hPSC-vSMC produced significantly larger forces than DMSO-treated hPSC-vSMC controls (p<0.005), adult AoSMC (p<0.0001) or fetal AoSMC (p<0.05). The AoSMC forces were measurably smaller than the DMSO controls (p<0.05), while the fetal AoSMC forces were not statistically different from the controls (Supplemental materials online, *Table S3*). MEKi treatment increased force generation in adult AoSMC, consistent with the Western blot data (*Figures 4E, 4F*), but not for AoSMC_f_ (Supplementary materials online, *Figure S12D)*.

#### 3.6.2 SMTs

We constructed humanized smooth muscle microtissues (SMTs), containing components only of human origin, from synthetic vSMCs in arrays of microfabricated tissue gauges (µTUGs) as illustrated schematically (*Figures 6A*,*6B)*. The microtissues developed tone (static force generation) as they compacted over ∼24-36 h. Representative traces of microtissue force vs time during compaction are shown in *Figure 6C*. A representative example of mixtures of vSMCs and ECM (human collagen and fibrinogen) seeded into each well following compaction to form a suspended microtissue structure spanning the pillars is shown in *Figure 6D*. Such microtissues could be maintained in culture for up to 6-8 days, and during this time, no “spontaneous” contractions with increased force development were observed.

**Figure 6.**
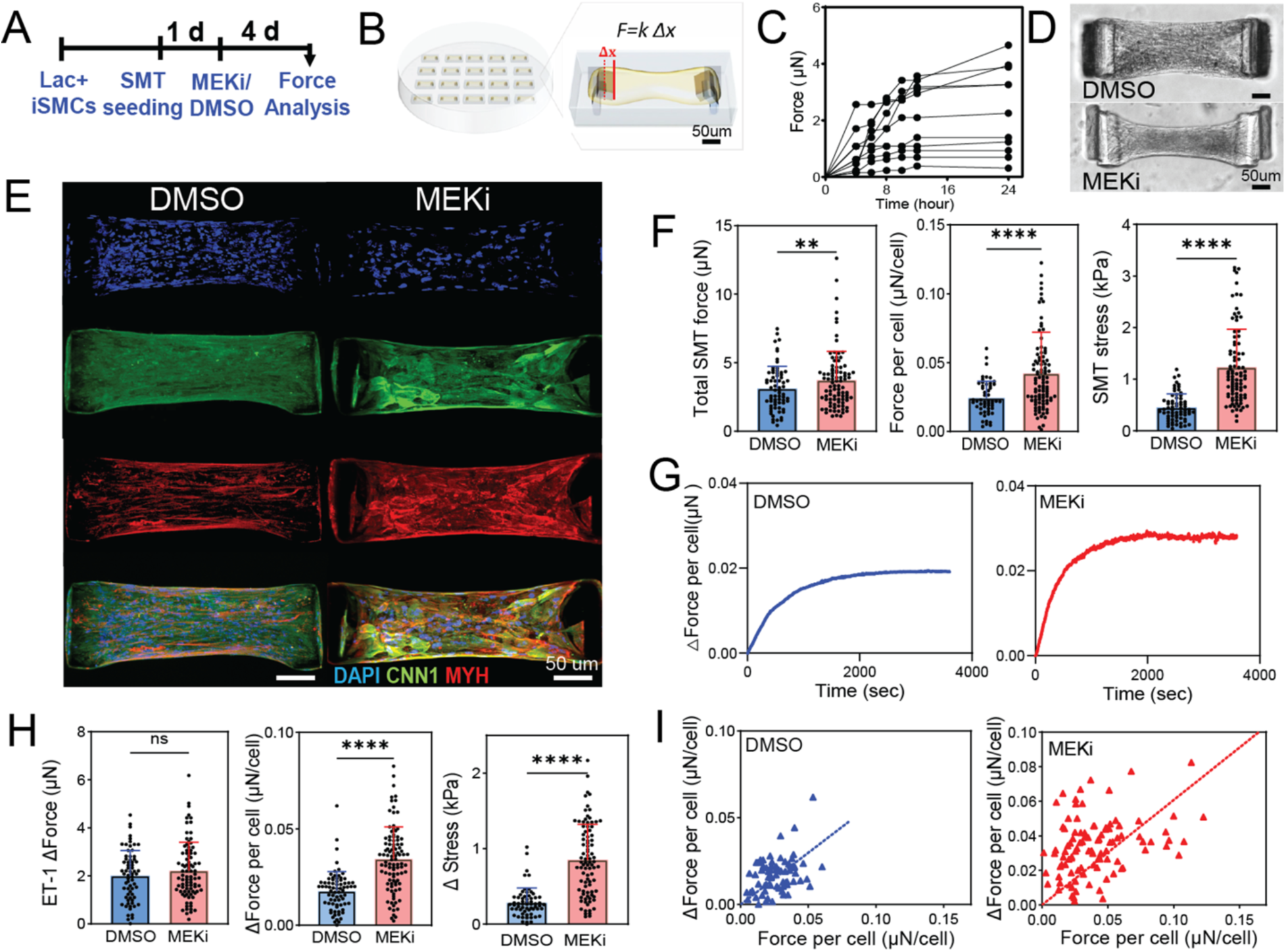
Force measurements of humanized microtissues containing JHU001-vSMC (SMT). Schematics illustrating A) the time-course of microtissue experiments, and B) a µTUG array and an expanded view of a single microtissue. Pillar deflection Δx reports the microtissue force with k = 0.25 µN/µm or 0.5 µN/µm. C) Representative traces of microtissue force vs time during tissue compaction. D) SMTs cultured after compaction in DMSO or with MEKi for 6 days. Scale bar 50 μm. E) SMT immunostaining showing the presence of CNN1 and non-specific MYH aligned along the long axis of the microtissues. Scale bar 50 μm. F) Microtissue force was 3.12±1.66 µN vs. 3.72±2.11 µN (*p*=0.0074); force per cell was 0.02±0.01 µN vs. 0.04 ± 0.03 µN (*p*<0.0001); and stress was 0.45±0.26 µN/µm² vs. 1.23±0.73 µN/µm² (*p*<0.0001), for SMTs treated with DMSO and MEKi obtained from µTUGs (k = 0.25 µN/µm). Error bars: ± SD; DMSO n=68, MEKi n=93. G) Exposure of SMTs to 1 µM ET-1, showing increased force generation with time. H) Changes in force generation following addition of ET-1 to SMT. Force was 2.00±1.05 µN in DMSO vs. 2.21±1.18 µN in MEKi (ns); Δ force per cell was 0.02±0.01 µN vs. 0.03±0.02 µN (*p*<0.0001); and Δ stress was 0.29±0.19 µN/µm² vs. 0.85±0.47 µN/µm² (*p*<0.0001). Error bars: ±SD; DMSO n=68, MEKi n=88. I) Correlations between ET-1 response and baseline SMT force. DMSO: slope = 0.59, R^2^=0.16, n=68. MEKi: slope = 0.52, R^2^=0.69, n=88. Statistics were performed using a Student’s *t-*test in F and H. *P*<0.05 was considered statistically significant (**p<0.01, ****p<0.0001).

To study contractile vSMC, SMTs were formed from synthetic JHU001- and WTC11-derived hPSC-vSMC and allowed to compact for 24 hrs, followed by treatment ± PD0325901 for 6 days (*Figure 6D*). The cells (JHU001) in these tissue constructs have strong signals for CNN1 and non-specific MYH, and are generally aligned along the long axis of the microtissues (*Figure 6E*). The developed force for these two conditions on µTUGs with *k =* 0.25 µN/µm is shown in *Figure 6F*. The MEKi treatment yielded a significant increase in total SMT force, force per cell, and force per cross-sectional area (SMT stress), consistent with the effects seen above in the single-cell force measurements with mPADs. Similar results were obtained for WTC11 and H9 SMTs (Supplementary materials online, *Figures S13, S14),* and corresponding results were obtained from JHU001- and WTC11-vSMC on µTUGs with stiffer pillars and *k =* 0.5 µN/µm (Supplementary materials online, *Figures S15A* and *S15E*).

To obtain a measure of their maximum force generation capability, we treated the SMTs with the vasoconstrictor endothelin-1 (ET-1). Exposure to 1 µm ET-1 led to increased force generation over timescales of ∼1,000 s, as illustrated in *Figure 6G* for JHU001 SMTs. Although the total change in force caused by ET-1 did not change significantly among the SMTs, larger increases were observed for the MEKi-treated JHU001 SMTs when expressed as force per cell or as a change in stress (*Figure 6H*). Similar results were found for WTC11 and H9 SMTs (Supplementary materials online, *Figures S13, S14*). The increases in force correlate with the baseline forces, as shown in *Figure 6I* with slopes of ∼0.5 for the JHU001 SMTs. The increase in force generation of WTC11-vSMC caused by ET-1 was less pronounced than in JHU001-vSMC, and the increases were less well correlated with baseline force for the WTC11 and H9 SMTs (Supplementary materials online, *Figure S13, S14*). Qualitatively similar results for the ET-1 response were obtained on µTUGs with stiffer pillars with *k =* 0.5 µN/µm (Supplementary materials online, *Figure S15 B-D and F-H*).

Microtissues formed from AoSMC, as described in the schematic (*Figure 7A*), showed compaction dynamics, high vSMC viability (Calcein AM staining, >90%), structure and force generation comparable to that observed in hPSC-vSMC. The AoSMC microtissues developed tone as they compacted over ∼24 h (*Figure 7B*) and the tissue constructs had strong signals for CNN1 and non-specific MYH (*Figure 7C*), which showed the tendency for cellular alignment along the SMTs’ long axis. The three measures of cell mechanics for the AoSMC SMTs, total SMT force, force per cell and stress, fell within the ranges observed for the hPSC SMTs, but substantial batch-to-batch variations were observed in force generation, in both the total SMT force and the more intensive measures of force per cell and stress *(Figure 7D*). The dynamic ET-1 response of the AoSMC again developed over 1000s timescales (*Figure 7E*), but the magnitude of the ET-1 response varied considerably among batches (*Figure 7F*), with only a rough correlation between the tissue’s baseline force and the magnitude of the ET-1 response (*Figure 7G*).

**Figure 7.**
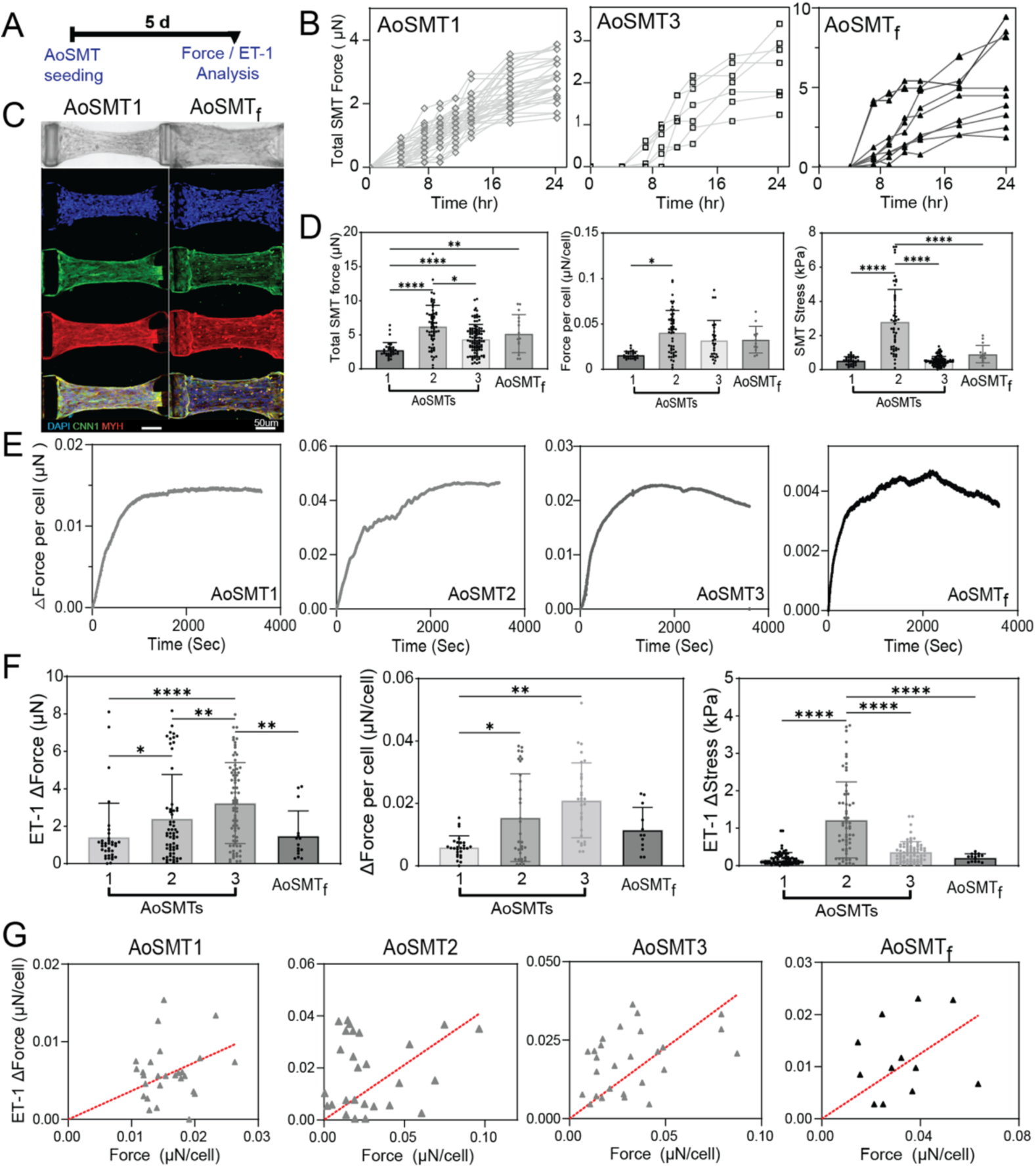
Force measurements of microtissues containing AoSMC (AoSMTs). A) Time course of SMT formation and measurements. B) Representative traces of microtissue force vs time as the tissues compacted for the indicated AoSMTs. C) White light images showing SMT structure and IF signals for CNN1 and MYH aligned along the long axis of the SMTs. Scale bar 50 μm. D) Microtissue force, force per cell, and stress for AoSMC SMTs. Error bars: ±SD. Total force and stress: AoSMC1 n=36, AoSMC2 n=66, AoSMC3 n=102, AoSMC_f_ n=15. Force per cell: AoSMC1 n=27, AoSMC2 n=51, AoSMC3 n=27, AoSMC_f_ n=12. E) SMT exposed to 1 µM ET-1, showing increases in force generation. F) Changes in force generation following addition of ET-1. Error bars: ±SD. Total force and stress: AoSMC1 n=36, AoSMC2 n=60, AoSMC3 n=75, AoSMC_f_ n=15. Force per cell: AoSMC1 n=27, AoSMC2 n=33, AoSMC3 n=30, AoSMC_f_ n=12. G) Correlations between ET-1 response and baseline SMT force. AoSMT1: slope = 0.36. R^2^=0.037, n=27. AoSMT2: slope = 0.42. R^2^=0.39, n=27. AoSMT3: slope = 0.45. R^2^=0.15, n=27. AoSMT_f_: slope = 0.31. R^2^=0.135, n=12. Statistics were performed using a two-way ANOVA in D and F. *P*<0.05 was considered statistically significant (*p<0.05, **p<0.01, ****p<0.0001).

## 4. Discussion

The use of hPSC to study human development and disease requires an understanding of how closely cell progeny are representative of somatic cells from tissues. Here, we investigated whether PM-derived hPSC-vSMC are comparable in phenotype and functional properties to commercially sourced human AoSMC. We assessed morphology, RNA and protein abundance, proliferative capacities, and we performed extensive analyses of the mechanics of single cells and of 3D tissues. Our findings underscore the significant heterogeneity inherent in AoSMC and validate the potential of hPSC-vSMC as a more consistent and reliable alternative for vascular biology and for tissue mechanic’s research.

An important observation from this study is the extensive heterogeneity of commercially sourced, proliferating AoSMC. Cells derived from aorta (mixed thoracic and abdominal) and cultivated in vitro under conditions identical to the hPSC-vSMC demonstrate prolonged doubling rates and differences in morphology, protein abundance, and mechanics. The use of Lac medium to enrich AoSMC proved to be equivocal. In one batch of AoSMC with a low purity of vSMC, short-term cultivation in Lac medium increased the MFI of ACTA2, CNN1 and TAGLN, but no changes in MFI could be demonstrated in the other batches of adult-derived AoSMC. Lac medium also increased transcripts encoding contractile vSMC markers in a batch-dependent manner. Reasons for these heterogeneous responses likely include proprietary but different expansion techniques of vSMC from donors of various ages, sex and genotypes. Heterogeneity among these primary lines could also be due to potential discrepancies among the regions of the aorta from which a given line was generated. Alternatively, the proliferating AoSMC (of unknown plasticity) may have been derived from isolated subsets of contractile vSMC undergoing phenotype switching within tissues.^29, 30^ Despite these variations, we found that MEKi can induce a more contractile phenotype in AoSMC, based on protein data and force measurements.

Enriched PM-derived hPSC-vSMC exhibit predictable and reproducible phenotypes, characterized by spindle-shaped morphologies at high density, a relatively low abundance of vSMC markers and an absence of MYH11 on Westerns. In contrast to published reports,^27^ our 2-day Lac enrichment protocol did not select for a more immature synthetic phenotype. No significant changes in RNA or protein levels for any SMC markers could be demonstrated relative to SMCM except for TAGLN, which has been reported to have moderate to high stability and to be regulated by mechanical tension.^31^ Instead, this protocol, originally used to enrich cardiomyocytes,^18, 32^ minimized the numbers of contaminating non-vSMC, overgrowth and cell death. Additionally, from cell surface capture proteomics data, the enriched cultures were negative for cardiac-(LSMEM2), endothelial-(CD31) and fibroblast-restricted (LCAT, LOXL2, LOXL4) markers,^33^ but positive for generic SMC markers (CD90^lo^, CD44^+^, CD13^+^, CD73^+^).^11^

Despite minor variations among batches and differences due to plating densities, PM-derived hPSC-vSMC exhibit phenotypic characteristics typical of synthetic cells. These in vitro differentiated vSMC proliferate, deposit ECM collagen, have low to undetectable levels of MYH11 and show density-dependent contact inhibition.^34^ Treatment with the MEK inhibitor, PD0325901 induced a significant shift toward a non-proliferating, contractile phenotype. The sizes of these cells, based on the literature, were larger than expected (possibly due to a transient loss of cell viability and cell spreading), but the phenotype was characterized by elevated quantities of contractile proteins and improved structural organization of actin filaments. Thus, the historical reliance on simple morphological characteristics to define vSMC phenotypes may be insufficient and potentially inaccurate. Upon MEKi withdrawal, these cells could switch back to a proliferating phenotype. These observations reinforce previous reports^11^ indicating that MEK inhibition is an effective approach for driving phenotype maturation in vitro. However, rapamycin did not induce a contractile phenotype switch in hiPSC-vSMC (Supplementary Material Online*, Figure S7*).^28, 35^ Thus, future studies will be required to assess how compounds like all-trans retinoic acid or RepSox affect hPSC-vSMC phenotype switching.^36, 37^

Our functional evaluation of normal synthetic and contractile hPSC-vSMC has established the baseline mechanical properties of these cells both in 2D and 3D. While other studies have evaluated cell traction force^38^ and contractility responses to agonists,^5, 39^ here we have provided a direct quantitative comparison of force generation by AoSMC and by PM-vSMC from hPSC. Using mPADs, we quantified single-cell contractile forces (analogous to low density cultures) and demonstrated that synthetic hPSC-vSMC generated forces comparable to those of human AoSMC. Notably, MEKi treatment significantly enhanced these forces, further aligning the functional properties of these cells with more mature, contractile AoSMC (MEKi-treated). The average magnitude of the force per post (*F_avg_*) produced by these cells on mPADs, in the range of ∼2 nN/post, is comparable to that observed on mPADs in vSMC from other sources, including bovine pulmonary artery SMC^40^ and porcine artery SMC,^41^ accounting for differences in post stiffness and spacing. The total average forces per cell are somewhat larger in these systems, likely due to the cells’ larger size. Overall, these quantitative validations establish a critical baseline foundation for utilizing hPSC-vSMC derived from paraxial mesoderm for disease modeling, drug testing platforms where functional readouts are essential, and for the analysis of genetic mutations that may affect vSMC function.

SMTs demonstrated responsiveness to contractile stimuli like ET-1. However, some differences in the measures of force per cell were observed between the SMT and the mPAD approaches. The SMTs measure forces generated along the cells’ polarization axis in a 3D environment with both cell-cell and cell matrix contacts. While it is tempting to seek a direct comparison of the SMT force per cell (∼30-50 nN/cell) to the single-cell results by scaling *F_avg_* (average magnitude of force per post) from the mPADs by the number of posts per cell (typically ∼100 for JHU001-vSMC), this value of ∼200 nN/cell overestimates the force applied to a neighboring body along the polarization axis by at least a factor of two,^41^ and likely more as it includes forces perpendicular to that axis.^40^ Disparities in force generation also can arise from differences in cellular organization resulting from the 2D vs 3D environments, as well as differences in effective substrate stiffness.^42^ The stiffness of mPADs is well-characterized (∼12 kPa),^43^ but for SMTs it can depend on details of the matrix organization, cell density, cell-cell and cell matrix interactions,^40, 44^ and thus it is likely that in-situ measurements of SMT stiffness would be required for detailed quantitative comparisons. The correspondence and correlations between the results in the 2D and 3D geometries provides important confirmation of the main observations of vSMC mechanics revealed by the current studies; however, the lack of additional data evaluating other known stimuli that may yield different responses in “immature” vs adult vSMC represents a functional limitation to the current study. An additional discussion of strengths and limitations can be found in Supplementary Material Online, Discussion.

In conclusion, our comprehensive analysis underscores the substantial variability among commercially sourced AoSMC. Experimentally, PM-derived hiPSC-vSMC offer a robust, reproducible, and functionally relevant alternative that can be precisely manipulated to model synthetic and contractile phenotypes. Notably, the mechanical properties of the AoSMC are well-recapitulated by the hiPSC-vSMC across multiple lines, validating this platform for the study of vSMC mechanics and how growth factors, gene mutations or pharmacological agents may affect cellular forces that might contribute to disease states. This study thus establishes hiPSC-vSMC as a viable and versatile platform for studying vSMC mechanics in a wide range of human vascular diseases, and the PM-cell type developed and profiled here represents an as yet underutilized in vitro model system for investigating genetic/inflammatory/metabolic drivers of aneurysms of the descending or abdominal aorta.

## References

1. Kwartler CS, Pinelo JEE. Use of iPSC-Derived Smooth Muscle Cells to Model Physiology and Pathology. Arterioscl Throm Vas 2024;44:1523–1536.

2. Elmarasi M, Elmakaty I, Elsayed B, Elsayed A, Al Zein J, Boudaka A, Eid AH. Phenotypic switching of vascular smooth muscle cells in atherosclerosis, hypertension, and aortic dissection. Journal of Cellular Physiology 2024;239:e31200.

3. Majesky MW. Developmental basis of vascular smooth muscle diversity. Arterioscl Throm Vas 2007;27:1248–1258.

4. Shen MC, Quertermous T, Fischbein MP, Wu JC. Generation of Vascular Smooth Muscle Cells From Induced Pluripotent Stem Cells Methods, Applications, and Considerations. Circ Res 2021;128:670–686.

5. Cheung C, Bernardo AS, Trotter MWB, Pedersen RA, Sinha S. Generation of human vascular smooth muscle subtypes provides insight into embryological origin-dependent disease susceptibility. Nature Biotechnology 2012;30:165–173.

6. MacFarlane EG, Parker SJ, Shin JY, Kang BE, Ziegler SG, Creamer TJ, Bagirzadeh R, Bedja D, Chen YC, Calderon JF, Weissler K, Frischmeyer-Guerrerio PA, Lindsay ME, Habashi JP, Dietz HC. Lineage-specific events underlie aortic root aneurysm pathogenesis in Loeys-Dietz syndrome. J Clin Invest 2019;129:659–675.

7. Hu YW, Cai ZH, He B. Smooth Muscle Heterogeneity and Plasticity in Health and Aortic Aneurysmal Disease. Int J Mol Sci 2023;24:11701.

8. Pfaltzgraff ER, Bader DM. Heterogeneity in Vascular Smooth Muscle Cell Embryonic Origin in Relation to Adult Structure, Physiology, and Disease. Dev Dyn 2015;244:410–416.

9. Bentzon JF, Majesky MW. Lineage tracking of origin and fate of smooth muscle cells in atherosclerosis. Cardiovasc Res 2018;114:492–500.

10. Bochaton-Piallat ML, Clowes AW, Clowes MM, Fischer JW, Redard M, Gabbiani F, Gabbiani G. Cultured arterial smooth muscle cells maintain distinct phenotypes when implanted into carotid artery. Arterioscl Throm Vas 2001;21:949–954.

11. Kumar A, D’Souza SS, Moskvin OV, Toh H, Wang BW, Zhang J, Swanson S, Guo LW, Thomson JA, Slukvin II. Specification and Diversification of Pericytes and Smooth Muscle Cells from Mesenchymoangioblasts. Cell Rep 2017;19:1902–1916.

12. Maguire EM, Xiao QZ, Xu QB. Differentiation and Application of Induced Pluripotent Stem Cell-Derived Vascular Smooth Muscle Cells. Arterioscl Throm Vas 2017;37:2026–2037.

13. Donadon M, Santoro MM. The origin and mechanisms of smooth muscle cell development in vertebrates. Development 2021;148:dev197384.

14. Wanjare M, Kuo F, Gerecht S. Derivation and maturation of synthetic and contractile vascular smooth muscle cells from human pluripotent stem cells. Cardiovasc Res 2013;97:321–330.

15. Chua C, DiSilvestre D, Joshi-Mukherjee R, Tung L, Tomaselli G, Boheler KR. Generation of an induced pluripotent stem cell line, JHUi008-A, from a healthy female donor. Stem Cell Research 2025;88:103819.

16. He J, Weng Z, Wu SCM, Boheler KR. Generation of Induced Pluripotent Stem Cells from Patients with COL3A1 Mutations and Differentiation to Smooth Muscle Cells for ECM-Surfaceome Analyses. In: Boheler K, Gundry R, eds. The Surfaceome. New York, NY: Humana Press, 2018:261–302.

17. Kilpinen H, Goncalves A, Leha A, Afzal V, Alasoo K, Ashford S, Bala S, Bensaddek D, Casale FP, Culley OJ, Danecek P, Faulconbridge A, Harrison PW, Kathuria A, McCarthy D, McCarthy SA, Meleckyte R, Memari Y, Moens N, Soares F, Mann A, Streeter I, Agu CA, Alderton A, Nelson R, Harper S, Patel M, White A, Patel SR, Clarke L, Halai R, Kirton CM, Kolb-Kokocinski A, Beales P, Birney E, Danovi D, Lamond AI, Ouwehand WH, Vallier L, Watt FM, Durbin R, Stegle O, Gaffney DJ. Common genetic variation drives molecular heterogeneity in human iPSCs Nature 2017;546:370–375.

18. Hawthorne RN, Blazeski A, Lowenthal J, Kannan S, Teuben R, DiSilvestre D, Morrissette-McAlmon J, Saffitz JE, Boheler KR, James CA, Chelko SP, Tomaselli G, Tung L. Altered Electrical, Biomolecular, and Immunologic Phenotypes in a Novel Patient-Derived Stem Cell Model of Desmoglein-2 Mutant ARVC. J Clin Med 2021;10:3061.

19. Morrissette-McAlmon J, Xu WR, Teuben R, Boheler KR, Tung LS. Adipocyte-mediated electrophysiological remodeling of human stem cell - derived cardiomyocytes. J Mol Cell Cardiol 2024;189:52–65.

20. Geng YX, Wang ZJ. Review of cellular mechanotransduction on micropost substrates. Med Biol Eng Comput 2016;54:249–271.

21. Fu JP, Wang YK, Yang MT, Desai RA, Yu XA, Liu ZJ, Chen CS. Mechanical regulation of cell function with geometrically modulated elastomeric substrates. Nat Methods 2010;7:733–736.

22. Shi Y, Sivarajan S, Xiang KM, Kostecki GM, Tung L, Crocker JC, Reich DH. Pervasive cytoquakes in the actomyosin cortex across cell types and substrate stiffness. Integrative Biology 2021;13:246–257.

23. Legant WR, Pathak A, Yang MT, Deshpande VS, McMeeking RM, Chen CS. Microfabricated tissue gauges to measure and manipulate forces from 3D microtissues. Proc Natl Acad Sci U S A 2009;106:10097–10102.

24. Bose P, Huang CY, Eyckmans J, Chen CS, Reich DH. Fabrication and mechanical properties measurements of 3D microtissues for the study of cell-matrix interactions Methods Mol Biol 2018;1722:303–328.

25. Shi Y, Sivarajan S, Crocker JC, Reich DH. Measuring cytoskeletal mechanical fluctuations and rheology with active micropost arrays. Current Protocols 2022;2:e433.

26. Hannan RT, Peirce SM, Barker TH. Fibroblasts: Diverse Cells Critical to Biomaterials Integration. ACS Biomater Sci Eng 2018;4:1223–1232.

27. Yang LB, Gao L, Nickel T, Yang J, Zhou JY, Gilbertsen A, Geng ZH, Johnson C, Young B, Henke C, Gourley GR, Zhang JY. Lactate Promotes Synthetic Phenotype in Vascular Smooth Muscle Cells. Circ Res 2017;121:1251–1262.

28. Martin KA, Merenick BL, Ding M, Fetalvero KM, Rzucidlo EM, Kozul CD, Brown DJ, Chiu HY, Shyu M, Drapeau BL, Wagner RJ, Powell RJ. Rapamycin promotes vascular smooth muscle cell differentiation through insulin receptor substrate-1/phosphatidylinositol 3-kinase/Akt2 feedback signaling. J Biol Chem 2007;282:36112–36120.

29. Worssam MD, Lambert J, Oc S, Taylor JCK, Taylor AL, Dobnikar L, Chappell J, Harman JL, Figg NL, Finigan A, Foote K, Uryga AK, Bennett MR, Spivakov M, Jorgensen HF. Cellular mechanisms of oligoclonal vascular smooth muscle cell expansion in cardiovascular disease. Cardiovasc Res 2023;119:1279–1294.

30. Worssam MD, Lambert J, Oc S, Taylor JCK, Taylor AL, Dobnikar L, Chappell J, Harman JL, Figg NL, Finigan A, Foote K, Uryga AK, Bennett MR, Spivakov M, Jorgensen HF. Cellular mechanisms of oligoclonal vascular smooth muscle cell expansion in cardiovascular disease (vol 119, pg 1279, 2023). Cardiovasc Res 2025;121:367–367.

31. Liu R, Hossain MM, Jin JP. The Expression and Degradation of SM22-Alpha/Transgelin are Regulated by Mechanical Tension in the Cytoskeleton. Biophys J 2017;112:435a–435a.

32. Huang CY, Maia-Joca RPM, Ong CS, Wilson I, DiSilvestre D, Tomaselli GF, Reich DH. Enhancement of human iPSC-derived cardiomyocyte maturation by chemical conditioning in a 3D environment. J Mol Cell Cardiol 2020;138:1–11.

33. Luecke L, Waas M, Littrell J, Wojtkiewicz M, Castro C, Burkovetskaya M, Schuette EN, Buchberger AR, Churko JM, Chalise U, Waknitz M, Konfrst S, Teuben R, Morrissette-McAlmon J, Mahr C, Anderson DR, Boheler KR, Gundry RL. Surfaceome mapping of primary human heart cells with CellSurfer uncovers cardiomyocyte surface protein LSMEM2 and proteome dynamics in failing hearts. Nat Cardiovasc Res 2023;2:76–+.

34. Sun YY, Qin SS, Cheng YH, Wang CY, Liu XJ, Liu Y, Zhang XL, Zhang W, Zhan JX, Shao S, Bian WH, Luo BH, Lu DF, Yang J, Wang CH, Zhang CX. MicroRNA expression profile and functional analysis reveal their roles in contact inhibition and its disruption switch of rat vascular smooth muscle cells. Acta Pharmacol Sin 2018;39:885–892.

35. Martin KA, Rzucidlo KM, Merenick BL, Fingar DC, Brown DJ, Wagner RJ, Powell RJ. The mTOR/p70 S6K1 pathway regulates vascular smooth muscle cell differentiation. Am J Physiol-Cell Physiol 2004;286:C507–C517.

36. Jung C, Han JW, Lee SJ, Kim KH, Oh JE, Bae S, Lee S, Nam YJ, Kim S, Dang C, Kim J, Chu N, Lee EJ, Yoon YS. Novel Directly Reprogrammed Smooth Muscle Cells Promote Vascular Regeneration as Microvascular Mural Cells. Circulation 2025;151:1076–1094.

37. Zhang J, McIntosh BE, Wang BW, Brown ME, Probasco MD, Webster S, Duffin B, Zhou Y, Guo LW, Burlingham WJ, Kent C, Ferris M, Thomson JA. A Human Pluripotent Stem Cell-Based Screen for Smooth Muscle Cell Differentiation and Maturation Identifies Inhibitors of Intimal Hyperplasia. Stem Cell Reports 2019;12:1269–1281.

38. Biel NM, Santostefano KE, DiVita BB, El Rouby N, Carrasquilla SD, Simmons C, Nakanishi M, Cooper-DeHoff RM, Johnson JA, Terada N. Vascular Smooth Muscle Cells from Hypertensive Patient-Derived Induced Pluripotent Stem Cells to Advance Hypertension Pharmacogenomics. Stem Cells Translational Medicine 2015;4:1380–1390.

39. Dash BC, Levi K, Schwan J, Luo J, Bartulos O, Wu HW, Qiu CH, Yi T, Ren YM, Campbell S, Rolle MW, Qyang YB. Tissue-Engineered Vascular Rings from Human iPSC-Derived Smooth Muscle Cells. Stem Cell Reports 2016;7:11–20.

40. Liu AS, Wang H, Copeland CR, Chen CS, Shenoy VB, Reich DH. Matrix viscoplasticity and its shielding by active mechanics in a microtissue model: in situ experiments and mathematical modeling. Scientific Reports 2016;6:33919.

41. Nagayama K, Adachi A, Matsumoto T. Heterogeneous response of traction force at focal adhesions of vascular smooth muscle cells subjected to macroscopic stretch on a micropillar substrate. J Biomech 2011;44:2699–2705.

42. Petit C, Yousefi AAK, Ben Moussa O, Michel JB, Guignandon A, Avril S. Regulation of SMC traction forces in human aortic thoracic aneurysms. Biomechanics and Modeling in Mechanobiology 2021;20:717–731.

43. Weng SN, Fu JP. Synergistic regulation of cell function by matrix rigidity and adhesive pattern. Biomaterials 2011;32:9584–9593.

44. Zhao R, Chen CS, Reich DH. Force-driven evolution of mesoscale structure in engineered 3D microtissues and the modulation of tissue stiffening. Biomaterials 2014;35:5056–5064.

