## Supplemental Information for "Phenotypes and Cellular Mechanics of Primary Human Aorta- and Pluripotent Stem Cell-derived Vascular Smooth Muscle Cells"

From

1.Department of Medical Laboratory Science and Biotechnology, National Cheng Kung University, Tainan, Taiwan

2. Department of Physics and Astronomy, Johns Hopkins University, Baltimore, MD, USA.

3. Department of Biomedical Engineering, Johns Hopkins University, Baltimore, MD, USA

4. Division of Cardiology, Department of Medicine, Johns Hopkins University, Baltimore, MD, USA

**Key words:** human induced pluripotent stem cells, vascular smooth muscle cells, cell mechanics, phenotype switching

#### Contents

| | |
|---|---|
| p. 2 | Supplemental Methods |
| p. 13 | Supplemental Results |
| p. 20 | Supplemental Discussion |
| p. 24 | References Cited |
| p. 26 | Supplemental Figures |
| p. 42 | Supplemental Tables |

### Supplemental Methods

The methods described in this section are detailed and meant to complement the methods provided in the main text of the manuscript.

**Human PSC-vSMC and primary human Aortic Smooth Muscle Cells (AoSMCs) cell culture**

In vitro derived vSMC from paraxial mesoderm, corresponding to cells from the descending aorta, were differentiated from human induced pluripotent stem cell (hiPSC) lines JHU001 (female)[1] and WTC11 (male GM25256, Coriell Institute) and from the embryonic stem cell (ESC) line H9 (female, WiCell) according to protocols from Cheung et al.[2] with modifications.[3] Briefly, as outlined in *Figure 1A*, plated human pluripotent stem cells (hPSC) were passaged with TrypLE and plated on geltrex-coated dishes for 24-36 hours (Day -1 to -1.5). When hPSC colonies of ~50-100 cells were observed, the medium was switched to CDM (49% IMDM, 49% Ham's F12 Nutrient Mix, 1% chemically defined lipid concentrate, 15 μg/mL optiferrin, 7 μg/mL recombinant human insulin, and 450 μM monothiol glycerol) containing polyvinyl alcohol (0.5 g in 500 mL CDM) to make CDM-PVA complete medium. CDM-PVA was supplemented with 0.2% fibroblast growth factor 2 (FGF2), 0.1% bone morphogenetic protein 4 (BMP4) and 0.1% phosphoinositide 3-kinase inhibitor LY294002 to initiate differentiation to mesoderm at Day 0 for 1.5 days. Subsequently, cells were fed with medium containing 0.2% FGF2 and 0.1% LY294002 for 3.5 additional days. Then cells were passaged and cultured for 13-14 days in medium containing 0.1% platelet-derived growth factor BB (PDGF-BB) and 0.1% transforming growth factor beta 1 (TGFβ1). After 18 days of differentiation, vSMC were transitioned to smooth muscle cell medium (SMCM, ScienCell) containing 2% fetal bovine serum (FBS), and when confluent, expanded and frozen, usually at passage 1 or 2. Thawed cells were cultivated in SMCM, and confluent cells enriched in glucose-deficient medium containing lactate. Two media were tested for vSMC enrichment. The first consisted of RPMI 1640 supplemented with B27 (no insulin), 4 mM lactate (L-lactate), 2 ng/mL human fibroblast growth factor 2 (FGF2) and 0.5 ng/mL human epidermal growth factor (hEGF) for 6 days i.e., Lactate + growth factors (Lac + GF).[4] The second contained DMEM supplemented with 4 mM lactate for 2-6 days (i.e., Lac).[5] Following enrichment, cells were switched back to SMCM and cultivated as monolayers for 1-4 days to allow recovery and the formation of confluent cells with a fusiform or spindle-like shape. After the conclusion of these tests, all subsequent enrichment experiments were performed using the Lac medium.

AoSMC were obtained from ATCC, Lonza and ScienCell. The cell batch numbers and characteristics are shown in Table 1; however, it is noteworthy that the suppliers did not indicate from which portion of the aorta the cells were obtained. It is likely that the cells came from both ascending and descending thoracic aorta and from the abdominal aorta. Cells were plated according to the manufacturer's instructions, followed by a stepwise transition to complete SMCM, so that all of the cells used in this study were cultured under the same conditions. All in vitro-derived vSMC and AoSMC were analyzed between passages 2 and 10.

#### Phenotypic Analyses of vSMC

##### *Cell Area, Aspect Ratio, and Nuclear Size Analysis*

Monolayer cultures of vSMC (hPSC- and AoSMC) were imaged using an EVOS inverted microscope. Cell area and aspect ratio were quantified from fluorescent images (see below) using ImageJ (NIH). For each analysis, individual cells were selected and cell boundaries were manually traced to generate contours. Nuclear size was measured separately by manually outlining nuclei on DAPI fluorescent images acquired with an EVOS M5000 Imaging System (ThermoFisher). The measured pixel values for both cell and nuclear areas were converted to micrometer values using calibration factors and data were tabulated in Excel for statistical analysis.

***RNA Assessments***

RNA was prepared using the RNeasy Plus Mini Kit (Qiagen, Hilden, Germany) according to the manufacturer's protocol, transferred to 1.5 mL tubes, and stored at −80 °C. The concentrations of isolated RNA were determined using a NanoDrop spectrophotometer (ThermoFisher). Reverse transcription was performed using the High-Capacity cDNA Reverse Transcription Kit for iPSC-derived cell lines and Super Script IV cDNA Synthesis Kit for the primary cell lines (ThermoFisher Scientific) with 0.5-1 µg of RNA in a 20 µL reaction volume according to the manufacturer's instructions. cDNA was quantified using the Qubit ssDNA Assay Kit (ThermoFisher Scientific), diluted to 10 ng/µL for hPSC-vSMC, and stored at −20 °C. Primary line cDNA was diluted to 2 ng/µL. The abundance of vSMC-restricted contractile proteins (calponin 1 - CNN1; smooth muscle actin - ACTA2; transgelin – TAGLN; smooth muscle myosin heavy chain - MYH11; smoothelin - SMTN), extracellular matrix proteins (collagen 1, subunit A1 - COL1A1; collagen 3, subunit a1 - COL3A1; elastin - ELN) and intermediate filament proteins (vimentin - VIM) were measured by quantitative PCR (qPCR). RNA encoding non-muscle myosin heavy chain isoforms 9 and 10 (MYH9, MYH10) were also assessed. The primer sets used for SYBR Green-based reactions are shown in Table S1. Quantitative PCR (qPCR) was conducted using the PowerTrack SYBR Green Master Mix (ThermoFisher) and assayed using the ViiA 7 Real-Time PCR System (Applied Biosystems). Each reaction was carried out in a 10 µL volume containing 5 µL of PowerTrack SYBR Green Master Mix, 0.5 µL of each forward and reverse primer, 2.5 µL of Molecular Biology Grade Water (Corning), and 2 µL of cDNA template. qPCR plates were robotically loaded on a Biomek 4000 (Beckman Coulter) and run in triplicate on all targets. Primary cell lines were run in duplicate. The qPCR data were processed and analyzed in R. Outliers were systematically determined and removed as CT values that varied from the mean of each triplicate by more than 2 units. Relative expression of targets was determined using the $2^{-\Delta\Delta Ct}$ method using RPL32 as an internal control for normalization to ensure accurate quantification. Comparisons between conditions were made using a Welch's *t*-test. In some experiments, qPCR analyses were expressed as $2^{-\Delta\Delta CT}$. In some qPCR experiments performed by one of the laboratories, probe-based real-time PCR techniques were used to quantify cDNA levels generated from RNA. Taqman data comparing vSMC treated with Lac medium versus SMCM are presented as a ratio of the $\log_2$ fold change. Relevant information regarding the forward and reverse primers and oligonucleotide-labelled probes for Taqman analyses are provided in Supplemental Table S1.

***Protein Analyses***

For immunostaining, cells were seeded into 6-well plates at a seeding density of 100,000 cells/mL. The cells were cultured overnight for subconfluent cultures or allowed to grow for ~5-7 days for confluent cultures, washed with DPBS and fixed for 10 minutes at room temperature in 4% paraformaldehyde. Wells were washed 3 times with DPBS and the cells were permeabilized with 0.1% Triton in PBS for 10 minutes. Fixed cells were incubated in blocking solution, consisting of 1% bovine serum albumin solution in DPBS for 1 h at room temperature. Primary antibodies against CNN1, ACTA2, MYH9, MYH10, MYH11 and TAGLN were added and incubated overnight on 4 °C at a dilution between 1:20 and 1:100 in blocking solution. Samples were washed three times using DPBS. Secondary antibodies (1:100; ThermoFisher Scientific),

DAPI (1:200; Thermo Fisher Scientific), and Phalloidin (1:200; ThermoFisher Scientific), were incubated for 1 hour at room temperature in the dark. Images were acquired using the Zeiss LSM 780 Confocal Laser Scanning Microscope (Carl Zeiss AG) and the corresponding software Zen (Carl Zeiss AG). Antibody information is provided in Table S2.

For Western blots, cells were kept on ice during protein extraction. After two washes with DPBS, cells were lysed in RIPA buffer (RIPA1, Cyrusbioscience) supplemented with protease inhibitor (1:200 dilution). Lysates were collected using a cell scraper, transferred to Eppendorf tubes, and either stored at –80 °C or processed immediately for protein quantification using the Qubit Protein Assay Kit (Q33211, ThermoFisher). For SDS-PAGE, equal amounts of protein (25 µg per sample) were separated on 4–12% NuPAGE Bis-Tris gels (NP0322, ThermoFisher) using Tris-MOPS-SDS Running Buffer (NP0001, ThermoFisher). Proteins were transferred to PVDF membranes (88518, ThermoFisher), which were blocked with 5% skim milk in 1× Tris buffered saline with 0.1% Tween 20 detergent for 90 min at room temperature. Blocked membranes were incubated with primary antibodies overnight at 4 °C (see Table S2 for complete antibody list, sources, and dilutions). Membranes were washed and incubated with appropriate HRP-conjugated secondary antibodies for 1 hr at room temperature. Protein bands were visualized using SuperSignal West Pico PLUS Chemiluminescent Substrate (34577, ThermoFisher), detected with an iBright imaging system and quantified using ImageJ software for image analysis. For total protein visualization, SDS-PAGE gels were stained with Coomassie Blue (PH1847, Phygene).

For flow cytometry, cells were fixed and permeabilized using the FIX & PERM Cell Fixation and Cell Permeabilization kit according to the manufacturer's protocol (ThermoFisher), followed by incubation with primary antibodies to detect expression of CNN1, MYH11, ACTA2, TAGLN and phalloidin, and subsequent incubation with secondary antibodies. Flow cytometry analysis was performed using CytoFLEX (Beckman Coulter) and analyzed using FlowJo software (FlowJo, LLC). A list of antibody information is provided in Table S2.

***Proliferation and Cell Cycle***

For the assessment of the number of cells in culture over time, hiPSC-derived vSMC were digested and passaged from confluent cultures into a 24-well plate, coated with gelatin, and allowed to attach for 24 hours. At day 0 (after 24+ hours of attachment), treatments with DMSO (vehicle control) and 1 μM MEKi were initiated and initial counting measurements made. To determine cell counts, medium was aspirated, the well washed once with 1 mL DPBS, and the cells treated with 0.5 mL with TrypLE for at least 10 minutes. TrypLE was inhibited by the addition of an equal volume (0.5 mL) of SMCM, containing 2% FBS. Cells were fully detached by trituration. 10 μL of the cell solution was added to each side of a hemocytometer and the number of cells counted manually and averaged. Samples were collected every two days through day 10.

DNA cell cycle analysis was measured in fixed cells using propidium iodide (10 mg PI, Sigma)-stained nuclei by Flow cytometry.[6, 7] Cells were trypsinized (both the floating and the adherent cells were collected) and fixed with ice cold 70% ethanol. Cell cycle compartments were deconvoluted from single-parameter DNA histograms of 10,000 cells. To block the vSMC in G2, Lac-enriched cells were treated with 100 ng/mL nocodazole (Noc, Sigma) for 24 to 48 hours.

***DNA Content – PicoGreen assays***

Monolayers of vSMC were lysed in DNA lysing solution (10 mM Tris (Millipore Sigma, USA), 1 mM EDTA (Millipore Sigma, USA, 0.1% Triton X-100 (Millipore Sigma, USA) containing 0.1 mg/mL proteinase K (Qiagen, USA); 500 μL). The lysate, after transferring to

Eppendorf tubes, was heated at 50 °C overnight to digest proteins and inactivate nucleases. Samples were subsequently frozen at -20 °C. To quantify DNA content, samples (100 µL) were incubated with an equal volume of 1X PicoGreen (200X stock) (ThermoFisher) reagent and transferred (total 200 µL) to a black, flat 96-well plate along with DNA standards according to the manufacturer's instructions. Fluorescence was measured at Ex485nm/Em528 nm using a fluorescent plate reader.[8]

***Cell Viability and Live Dead Cell Staining***

Cell viability was assessed using the PrestoBlue Cell Viability Reagent (ThermoFisher) or Calcein AM (ThermoFisher). Ten volumes of pre-warmed expansion medium were mixed with one volume of PrestoBlue, and live cells in 96-well plates were incubated with this mixture for 1 hour at 37 °C in 5% $CO_2$. Absorbance at 590 nm was measured on an Infinite M200 PRO plate reader (Tecan), and blank values were subtracted. Live/dead staining was performed with the LIVE/DEAD™ Cell Imaging Kit (488/570) (ThermoFisher) according to the manufacturer's instructions. Cells plated in 24-well plates were incubated at room temperature for 15 minutes with equal volumes of the Live (green) and Dead (red) reagents. Images were acquired on an EVOS M5000 Imaging System (ThermoFisher).

**Force Measurements**

***Micropost Array Detector (mPAD) system***

For the quantification of the contractile forces of individual cells, micropost array detector (mPAD) devices were fabricated by replica molding.[9, 10] Poly(dimethylsiloxane) (PDMS) negatives mixed 10:1 with crosslinking agent were formed from Si originals and treated overnight with silane gas before use, following established techniques. PDMS arrays of microposts were then cast from the negatives. The cylindrical microposts had a diameter of 1.8 μm and a height of 6.4 μm, spaced by a lattice constant of 4 μm on a hexagonal lattice. Under small lateral deflections, each micropost had an effective spring constant $k$ = 15.7 nN/μm.[10]

Devices were fabricated by treating a glass coverslip with UV light and ozone using a Jelight UVO-cleaner for 7 minutes, then adding a drop of degassed 10:1 PDMS, stamping the negative mold onto this drop, then curing at 70˚C overnight. Once the devices were cured solid, the interface with the stamp was submerged in 70% ethanol and the stamp was peeled away from the device. The devices on the glass slides were then dried in a critical point dryer (Tousimis Samdri-795).

To functionalize these devices to ensure good contact with cells, blocks of cured PDMS mixed 1:25 and cut to be just larger than the face of an mPAD were wetted with 100 μL of 50 μg/mL fibronectin solution spread across the entire surface of each block and allowed to adsorb for one hour. After this adsorption period, dry mPADs were treated with UV/Ozone for 7 minutes and the fibronectin-treated blocks were submerged in DI water, dried with nitrogen gas, then stamped onto a dry mPAD with the treated face and pressed gently to ensure contact between the pillars and fibronectin with minimal damage. The interface between the device and stamp was submerged in 70% ethanol and the stamps were knocked off the devices very gently. The stamped mPADs were alternated between baths of 100% ethanol and DI water twice, then placed in 0.2% Pluronics F-127 (P6866, ThermoFisher) solution, where they were treated for 20 minutes. The functionalized devices were rinsed twice in a bath of DI water and stored in a bath of DPBS.

To seed vSMC onto functionalized devices, cells in culture were passaged as usual: using one rinse with DPBS, then an incubation with TrypLE express solution, followed by trypsin neutralization and cell collection with SMCM, centrifugation, aspiration of the supernatant, and resuspension in SMCM. To maximize the placement of cells directly onto the device, mPADs were placed in a 35 mm diameter well with SMCM before cells were resuspended in 0.5-3 mL of medium, depending on density in culture, and added dropwise directly onto the device. Devices were moved gently to an incubator and maintained for 24 hours to allow time for cell adhesion.

To collect data that demonstrates the force a single cell exerts on the pillars of an mPAD, each device was imaged using brightfield microscopy at 40x magnification on a Nikon TE-2000E microscope. Proper alignment for data acquisition was ensured by bringing the focal plane to the tops of the pillars on the mPAD. Videos were recorded for 1-10 seconds of isolated cells on an mPAD using a Prosilica GX camera at a rate of 10 fps, following established techniques.[11]

***Humanized Smooth Muscle Microtissues***

Microfabricated tissue gauges (µTUGs) were fabricated by replica molding.[12, 13] Each microtissue well measured 800 µm x 400 µm x 125 µm (length x width x depth), and the pillars were spaced by 500 µm. The µTUGs treated with 0.2% Pluronic F-127 for 15 minutes were rinsed with DPBS and dried. An extracellular matrix (ECM) solution was prepared from the following components: M199 (11825015, Gibco), HEPES (15630-80, Gibco), $NaHCO_3$ (5% *w/v*) (S5761, Sigma-Aldrich), human fibrinogen (0.75 mg/mL) (F3879, Sigma-Aldrich), and human type I collagen (3.2 mg/mL) (5007, Advanced Biomatrix), and neutralized with NaOH (S2770, Sigma-Aldrich). The ECM solution was added to µTUG devices placed on ice packs (432014, Corning) and centrifuged for 2 minutes at 935 g and –9 °C.

Synthetic hPSC-vSMC or AoSMC cultured in SMCM were suspended in ECM solution, loaded into µTUGs, and spun down into the microwells at 234 *g* for 2 minutes, with the device rotated 90° after 1 minute. Excess ECM solution was removed from every part of the device except the microtissue wells. Devices were centrifuged upside-down at 700 rpm for 20 seconds, rotating 90° after 10 seconds.[13] The devices were placed in an incubator for 25 minutes to allow the ECM to solidify within the microwells. SMCM was then added, and the devices were incubated for 24 h at 37 °C and 5% $CO_2$. After 24 h, medium was refreshed for control (SMCM supplemented with 1:1000 DMSO) or switched to SMCM containing MEKi for contractile induction. Cultures were maintained for an additional 4 days, except for those used to monitor cell compaction.

Contractile forces generated by SMTs were quantified from micropillar deflections in μTUG devices. Bright-field images of tissues and pillars were acquired at defined time points using an inverted microscope (Nikon TE-2000E or Olympus CKX-453). Pillar deflections were measured relative to their resting (undeflected) position using ImageJ/Fiji software, and forces were calculated by multiplying the deflection distance by the known spring constant of the micropillars (k = 0.25 or 0.5 μN/μm). To count cells for measurements of force per cell, the SMTs were stained with DAPI. When tissues appeared densely packed, multiple z-stack image fields (typically three per SMT, sampled across the tissue) were acquired. Nuclei were counted manually to avoid duplicate counting across overlapping regions. Cross-sectional areas of SMTs were estimated from Z-scan confocal images (Figure S16). A correlation parameter between tissue width and height was derived and applied to width measurements to calculate cross-sectional area. This enabled conversion of contractile force values into stress.[14] which reduces impacts due to size variability of the SMTs compared to measurements of the tissues' total force. Although this normalization reduces the impact of size variability, differences in tissue geometry remain an inherent limitation of microtissue assays. In addition, immunostaining in thicker SMTs may be affected by limited antibody penetration, which can produce stronger signal at the tissue periphery.

For the endothelin-1 (ET-1) response assay, SMTs were incubated in Tyrode's solution (in mM: 1.8 $CaCl_2$, 137 NaCl, 5.4 KCl, 15 glucose/dextrose, 1.3 $MgSO_4$, 1.2 $NaH_2PO_4$, and 20 HEPES; pH adjusted to 7.4 with NaOH) for 45 min to reduce basal vasoconstriction from serum-containing medium. Following equilibration, tissues were treated with 10 nM ET-1 and videos recorded at 1 frame/s for 60 min. Pillar deflections were quantified frame by frame, contractile force was calculated using the spring constant of the micropillars, and values were normalized to cross-sectional area to yield stress (*see Figure S15*). This allowed dynamic assessment of ET-1–induced contractile responses.

**Data Analysis**

***qPCR data analysis***

A Linear Mixed Model (LMM) was applied to normalize qPCR data across plates of hiPSC-vSMC by treating each plate as a random effect in the model to account for potential plate-to-plate variability (lmer(Mean_CT ~ 1 + (1|Plate)). To analyze the effects of the lactate or MEKi treatments on the cells, the relative expression of target genes was compared between treated and untreated cells (separated by cell line). Significance was determined using a Welch's (unequal variance) *t*-test (because the data failed to meet the assumptions of a Student's *t*-test). A summary of replicates is included under each plot.

***mPAD data analysis***

To analyze the acquired mPAD data, videos were processed using an algorithm[11] in Igor Pro 9 (WaveMetrics) which calculates the displacement of each of the posts beneath the cell from its predicted resting position. Using the effective spring constant k =15.7 nN/μm for each post, the force (deflection) exerted by a cell on each post was calculated, and the average force per affected post reported as a measure of total contractility for a single cell. As studies of MEKi treatment of primary AoSMCs (*Figure S12D)* showed large deflections of selected microposts near the periphery of the cells that were not adequately captured by our automated analysis routines, we updated this software to allow for manual determination of the deflections of such highly-deflected microposts as needed. Contraction force analysis code is available upon request.

***Statistics***

Plots, boxplots and scatterplots were constructed using GraphPad Prism7 (GraphPad. La Jolla, CA, USA). Data are presented as the mean ± standard deviation or as the mean ± standard error (SEM) with a corresponding n value. Statistical tests included a one-way or two-way ANOVA and a Student's *t*-test (two-sample, unpaired). Differences were considered statistically significant at $p<0.05$, and the level of significance is explicitly detailed either in the text or in the figures. All experiments were performed with an n = 3-7 replicates (as described in the text). Correlation was determined using linear regression.

#### Supplemental Results

##### Pilot Experiments for Human Pluripotent Stem Cell derived – vSMC Enrichment

Human pluripotent (induced and embryonic) stem cells (hPSCs) were cultured and differentiated to paraxial mesoderm derived vSMC as described.[2,3] Brightfield imaging of sub-confluent cells (1-4 days after passaging) had a mixed fusiform-like, rhomboid-like or fibroblast-like morphology (*Figure S1A*, DD18). At confluency, the cells developed an elongated, fusiform or spindle-shaped morphology (*Figure S1B*), but a few larger, fibroblast-like cells were observed that may have included some non-vSMC. Because glucose-free, lactate-containing medium has been reported to enrich for populations of muscle cells and to select for more synthetic vSMC,[4] we incubated the in vitro differentiated cells at >90% confluency with either SMCM medium or RPMI medium supplemented with lactate, hFGF2 and hEGF (Lac + GF) for 6 days[4] in a series of pilot experiments. VSMC cultured in SMCM were examined in parallel and used as controls. The results of these pilot experiments are provided as supplemental information to the main Methods and Results sections of this manuscript.

In these experiments, the control cells were spindle-shaped with a compact confluency (*Figure S1B*), but within 6 days, the cells often grew into layers. In Lac + GF enriched cells, we visually observed two main "phenotypes", which likely reflected the quality of the in vitro differentiations and ratio of vSMC to non-vSMC. In cases of sub-optimal differentiation (excluded from subsequent analyses), the population density of cells treated with Lac + GF decreased and became sub-confluent, likely due to the loss of relatively high numbers of non-vSMC from the culture (*Figure S1B*, 6 days). Post-enrichment, these cells usually needed an additional 3-7 days to become confluent. In the second, more common, case where the proportion of vSMC to non-vSMC was likely high, we observed an increase in vSMC densities as a function of time, i.e., the cells were growing in multiple layers upon other cells, similar to what we saw with the control cells grown in SMCM (*Figure S1B*, 6 days, inset).

To develop conditions where the cells were not overly confluent, we also tested vSMC enriched with DMEM medium lacking glucose, supplemented with lactate, but lacking any growth factors (Lac) as we had previously described for cardiomyocytes.[15] When cultured for 4-6 days in this medium, we observed significant cell death of both non-vSMC and vSMC (not shown).

However, when we incubated the cells for only 2 days in this medium, most rhomboid-shaped cells remained attached to the plate (*Figure S2C*). At this time, we observed both sub-confluent cultures, due to a significant loss of presumably non-vSMC, and relatively confluent cultures. Anecdotally, when sub-confluent enriched cells were switched back to SMCM after Lac-mediated cell enrichment, it took several days for the cultures to obtain a compact confluent state. If these cells were then replated, expanded and selected a second time with Lac, the confluency remained compact, suggesting that the original enrichment protocol had eliminated many or most of the non-vSMC. In the case where the cultures maintained a relatively high cell density post-enrichment, compact cell confluency and spindle-shaped cells were usually observed within 1-2 days.

#### Cell Phenotypes of Control and Enriched vSMC

##### *Immunostaining and Flow Cytometry*

The in vitro differentiated vSMC, irrespective of lactate enrichment, showed a high degree of fluorescent signal heterogeneity among contractile protein markers that we suggest is due to differences among cells cultured at high densities and not due only to poor penetration of antibodies into cells at confluency (based on replicate staining and flow cytometry). Only a small proportion (<1-5%) of DAPI-positive cells cultivated in SMCM lacked a signal for smooth muscle contractile proteins examined, excluding phalloidin (PHA), which was present in all cells (*Figure S2*, white arrows, Merge column). Although contractile-associated protein staining was mostly cytosolic, the signal intensities varied from cell to cell. This was especially pronounced for CNN1 (*Figure S2A*). TAGLN- and ACTA2-positive cells also showed signal variability; however, the signal intensities were similar, i.e., in cells where TAGLN was strongly stained, ACTA2 also showed strong signal intensities (*Figure S2B*). Interestingly, strong signals for PHA were also observed in cells that had weak signals for ACTA2 or TAGLN (*Figure S2B*). ACTA2 and TAGLN proteins, and to a lesser extent non-specific MYH11, also formed fiber-like structures in some of the plated cells, suggestive of a more mature phenotype. Although some of the observed variations might be due to the focal plane of the images, Z-stacks confirmed the variability of signal intensities among cells.

vSMC enriched in Lac + GF medium had the smallest size relative to either SMCM controls or Lac cultured cells (*Figure S2*). We attribute the compact size to the plating density of these cells, following 6 days of enrichment versus the 2 days of cultivation in SMCM or in Lac. Larger cells present in these cultures usually had proteins that formed fiber-like structures (*Figure S2B*, marked by white stars). DAPI-positive cells negative for contractile protein markers were rare, confirming that vSMC could be enriched by Lac + GFs. As with the untreated vSMC, cells were generally positive for contractile protein markers. Non-specific MYH11 and PHA had the most uniform signals (*Figure S2*), while ACTA2, CNN1 and TAGLN staining varied. In some cells, strong immunofluorescence staining for TAGLN was observed in cells weakly stained for ACTA2 and vice versa (*Figure S2B*), a finding that differed relative to SMCM cultivated cells.

When we examined vSMC following 2 days of cultivation in Lac medium with 1-2 days of recovery in SMCM, the culture density was generally lower than what we observed with either of the other two other cultivation conditions tested (*Figure S2*). Correspondingly, larger cells with a more rhomboid shape were observed more frequently; however, mostly spindle-shaped cells were present within 1-2 additional cultivation days. Nearly all the DAPI-positive cells were positive for at least one of the SMC contractile protein markers (TAGLN, CNN1, ACTA2, non-specific MYH11) (*Figure S2A*), consistent with vSMC enrichment. PHA staining was generally uniform among the cells. The signals for the other contractile proteins, and particularly CNN1, were not uniform from cell to cell. Many of the proteins also seemed to form fiber-like structures, similar to what we had observed under the other cultivation conditions (*Figure S2*). ACTA2 also had a more pronounced peri-nuclear staining pattern compared to the other two conditions. In some co-stained cells, fluorescent signals for TAGLN were greater than those observed for ACTA2, and signals for non-specific MYH11 were occasionally much higher than what we observed for CNN1 (*Figure S2A*). Non-specific MYH11 also seemed to have more fiber-like structures in these cells comparted to Lac + GF cultivated cells; however, this staining pattern may have been a function of cell density and size.

By qPCR, we examined transcripts encoding proteins associated with the extracellular matrix (ECM), intermediate filaments, and smooth muscle contraction (*Figure S3*). In these experiments, control vSMC cultured in SMCM generally had the lowest relative amounts of all of these transcripts when compared to either of the two lactate-treated cultures, possibly due to the presence of contaminating non-vSMC. Furthermore, we could not demonstrate any significant differences for any of the ECM transcripts analyzed in JHU001 (or WTC11) hPSC-vSMC between Lac + GF (6 days + 1-2 day recovery) and Lac (2 days + 1-2 days recovery) or between either lactate treatment compared to SMCM cultivated vSMC (*Figure S3C, D*, $p>0.05$). In JHU001 hPSC-vSMC, VIM was significantly decreased following enrichment in Lac + GF relative to Lac treated JHU001-derived vSMC, but the abundance of this transcript did not significantly differ between cells with Lac treatment and SMCM controls. In contrast, a significant increase in CNN1 and TAGLN transcripts was observed in Lac + GF-treated JHU001, but not WTC11, hPSC-vSMC, relative to SMCM controls. TAGLN was also increased in JHU001 hPSC-vSMC enriched with Lac relative to SMCM controls. In the WTC11 hPSC-vSMC, MYH11 RNA content was increased in the Lac + GF treated cells relative to both the Lac and the SMCM control groups (*Figure S3D*).

In Figure S3E, qPCR results performed with Taqman probes on line H9 ESC-vSMC are shown comparing the ratio of RNA levels between cells with Lac and SMCM medium.

Protein assessments were determined using Western blotting and flow cytometry. Western data are reported in the main text of the manuscript; however, images of the Westerns are provided from independent lysates of JHU001- and WTC11-vSMC (Lac vs SMCM)(*Figure S4A*). Data were normalized to total protein loading (*Figure S4B*), and summary data are provided (*Figure S4C*). Flow cytometry data are shown in Figure S5. Data were analyzed based on the gated region shown in the scatterplot, and histograms show data compared to unlabeled cells and isotype controls (*Figure S5A*). When we analyzed the fluorescent signals for $CNN1^{+}$, $ACTA2^{+}$, $TAGLN^{+}$ or non-specific $MYH11^{+}$ by flow cytometry (n=3, JHU001-vSMC and WTC11-vSMC), we did not observe any significant differences for any of the contractile proteins among cell populations cultivated with SMCM or enriched by either Lac + GF or Lac (*Figure S5*). The number of positive cells for all the markers varied between 90-95%, except for non-specific MYH11 in WTC11 cells, which varied between 45-55% (*not shown*). Analysis of the mean fluorescence intensity (MFI) of JHU001-vSMC showed that the signal intensities of ACTA2 and TAGLN were increased, although not significantly, in Lac + GF and Lac relative to SMCM cells (Lac + GF vs SMCM: or ACTA2: $p = 0.13$; TAGLN: $p=0.36$; Lac vs SMCM: ACTA2: $p = 0.33$; TAGLN: $p=0.20$, n=3, one-way ANOVA). Similar trends were observed in WTC11-vSMC. No significant difference could be demonstrated in the MFI for either CNN1 or non-specific MYH11 between cells cultured in SMCM or any of the enriched cells. These data are consistent with the immunostaining results described earlier, which showed some differences in signal intensity among a broad population of cells that were generally positive for at least one of the SMC-restricted markers. Data showing the relative levels of MYH9, 10, and 11 transcripts (JHU001) and Westerns (JHU001, H9) are described in the main text, and additional blots are shown in Figure S6.

Based on these data (IF, flow cytometry and qPCR) and the absence of MYH11 transcripts in DMSO treated vSMC, we conclude that treatment of vSMC by 4 mM Lac relative to SMCM controls yielded enriched, MYH11-negative vSMC that were largely devoid of non-vSMC .

***Effects of Rapamycin (Rap) on the phenotype of vSMC***

Kumar et al. reported that MEKi could induce immature, mesenchymoangioblast-derived synthetic vSMC (i.e., splanchnic mesoderm) to form a more mature, proliferatively quiescent, contractile phenotype.[16] As part of our tests to determine whether other small molecules could induce PM-originating vSMC derived from hPSC to develop a contractile phenotype, we treated cells with rapamycin (Rap), which has been implicated in the formation of a contractile phenotype in rodents, to induce a contractile phenotype.[17] Lac-enriched vSMC were treated every two days for 6-8 days with Rap and allowed to recover in SMCM for 1-2 days. DMSO was used as a vehicle control. Under control conditions, the enriched vSMC appeared as highly dense cultures, with spindle-shaped cells that were overly confluent (*Figure 2A*). Following 6 days of Rap treatment, at concentrations ranging from 0.1 to 20 nM, cells remained spindle-shaped, and the densities increased to a point where the vSMC were overly confluent (*Figure S7*), as was observed with control vSMC. No significant differences in DNA content could be demonstrated between Rap and DMSO treated cells; however, a decrease in total cell numbers was observed (<10% difference, data not shown). Because we did not see any significant difference in cell proliferation, cell numbers relative to controls, or in force generation relative to controls (not shown), no additional experiments are reported with Rap.

#### Supplemental Discussion

##### *Strengths and Limitations of Study*

As a strength of this study, we adapted an existing protocol[2, 3, 18] to generate vSMC from three hPSC lines. Use of multiple lines minimized potential qualitative and quantitative differences associated with interline variability, which can account for 5-46% of the variation in hiPSC phenotypes, including differentiation capacity and cellular morphology.[19] Moreover, the analysis of vSMC from multiple hPSC lines minimized possible epigenetic differences associated with inappropriate X-chromosome inactivation among female-derived lines (male: WTC11; female: H9, JHU001).[20, 21] Furthermore, we systematically examined the characteristics and mechanics of paraxial-mesoderm (PM)-derived vSMC from these hPSC lines, and we compared these profiles with aorta-derived primary vSMC purchased commercially. This is noteworthy, because vSMC from the PM lineage, which have been implicated in the development of aneurysms and other vascular lesions,[22, 23] have not been as thoroughly evaluated as vSMC derived from either the second heart field or neural crest.

In terms of hPSC-vSMC and AoSMC phenotypes, considerable effort was expended to quantify a number of cellular variables (e.g., cell size, proliferation, contact inhibition and aspect ratios). During these analyses, we determined that cell morphologies were dynamic and that vSMC shapes depended on plating density. At high densities, synthetic hPSC-vSMC assumed the "typical" spindle shape reported in the literature for contractile vSMC, but at low densities, these cells had the irregular, epitheloid-like phenotypes "typical" of synthetic vSMC.[24] These proliferating, but contact inhibited cells expressed a number of markers typical of this phenotype (e.g., COL1A1), while MEKi treatment induced a non-proliferative phenotype with enhanced expression of contractile markers (e.g., CNN1, ACTA2, MYH11). Moreover, the morphology of contractile hPSC-vSMC were not fully consistent with reported norms, as these cells were generally larger and less spindle shaped than anticipated.[24] It is currently unclear whether this morphological difference from the "normal" contractile vSMC shape is related to the in vitro differentiation of hiPSC, cell density or to the MEKi treatment.

A particularly noteworthy strength of the present study was the application of complementary biophysical approaches including quantitative 3D force measurements, analyzing over 60 microtissues per hPSC line, complemented by single-cell traction force measurements of individual cells on micropillars. Additional strengths include the mechanical characterizations and comparisons with primary human aortic SMCs (AoSMCs). The micropost assays evaluated single-cell force generation, while the 3D microtissue system reflected collective, tissue-level mechanics. The general agreement in data between these two approaches confirms the robustness of the mechanical phenotype, and the results allowed us to determine that the vSMC in 3D constructs did not show, at least in this in vitro setting, any cellular cooperative effects. The results also demonstrated that synthetic and contractile (MEKi-treated) hPSC-vSMCs can be reproducibly generated in vitro with proliferation and mechanical properties comparable to those produced by AoSMCs.

However, limitations do arise when attempting to systematically measure the contractile forces of isolated cells. As a case in point, our mPADs measurements were performed on those cells that adhered to mPADs coated with fibronectin. This coating differs from the plating conditions (e.g., gelatin) we used for all other measurements and culture conditions. Furthermore, we cannot account for trends in force generation that may correlate with fibronectin adhesion. We also determined the mechanics of cells that could be analyzed based on a designated set of imaging conditions; however, we also observed a few hPSC-vSMC that were elongated and very large (approximately 5x larger than average) following MEKi treatment that were difficult to assess. Although these cells were uncommon, our inability to adequately measure the forces produced by these cells likely caused us to underestimate the contractile behavior of the overall MEKi-treated vSMC population. In SMT (3D) cultures containing contractile vSMC within a microtissue structure, we also experienced some difficulty measuring the behavior of highly contractile tissues. Microtissues with either larger numbers of cells following compaction or containing cells that may contract more strongly could "peel off" the pillars of the device within a few days of formation. This may have caused us to underestimate the average forces generated by these tissues, as we cannot measure tissues which have come off the device.

As additional limitations, we primarily examined markers associated with the synthetic and contractile phenotypes of vSMC. However, vSMC possess high plasticity, and other phenotypes have been described in vascular aging, atherosclerosis, aortic aneurysm, and disease. These vSMC sub-types typically down-regulate contractile proteins such as ACTA2 and CNN1, and assume some cellular functions typical of osteoblast, fibroblast, macrophage, mesenchymal cells, as well as degradative vSMC.[24] This highly plastic phenotype transformation is regulated by a complex network consisting of circulating plasma substances, transcription factors, growth factors, inflammatory factors, non-coding RNAs, integrin family, and Notch pathway – none of which were anticipated to be prominent in the present in vitro study. Second, we focused on the role of MEKi to induce a more contractile phenotype. When we tested rapamycin, which in rodents had been reported to promote a contractile phenotype, we did not see a phenotype switch. These findings suggested that the PM-vSMC from human may have unique characteristics that differ from those in rodents. Additional studies with RepSox,[25] a small-molecule inhibitor of the TGF-β receptor that drives the differentiation of stem cells into pro-contractile vSMC, or with retinoic acid, which promotes fibroblast reprogramming into vSMC, would likely be needed to further validate possible phenotypic differences of contractile vSMC among animal models and humans. Furthermore, the contractile studies evaluating agonists were not fully exploited. We focused on endothelin-1 (ET-1) because it is a well-established, potent agonist widely used to evaluate smooth muscle contractility, and it provides a robust and reproducible readout across SMC models. We also performed a few experiments with phenylephrine, a selective alpha-1 adrenergic receptor agonist in vSMC. In these experiments (n=2-3), it produced contractile responses qualitatively similar to those observed with ET-1, but with smaller amplitudes. Despite these similarities, a systematic pharmacological analysis and comparisons of both pro-contractile and pro-relaxation agonists would be required to determine whether the contractile properties are fully conserved between hPSC-vSMC and AoSMC or whether some developmental or functional differences exist.

Finally, our focus on PM- vSMCs (versus other embryological origins) has specific benefits and applications for future studies. As described in the Introduction of this paper, aneurysm etiology caused by genetic defects depends on the lineage of origin and likely differs between ascending and descending/abdominal aorta. Hereditary factors usually dominate the former, while inflammatory and metabolic drivers are generally thought to dominate the latter. The PM-cell type developed and profiled here, thus, represents an underexploited in vitro model system for evaluating genetic, inflammatory, and metabolic drivers of aneurysms of the descending/abdominal aorta that may be implicated in the formation of vascular lesions in adults.

### References Cited

1. Chua C, DiSilvestre D, Joshi-Mukherjee R, Tung L, Tomaselli G, Boheler KR. Generation of an induced pluripotent stem cell line, JHUi008-A, from a healthy female donor. *Stem Cell Research* 2025;**88**:103819.
2. Cheung C, Bernardo AS, Trotter MWB, Pedersen RA, Sinha S. Generation of human vascular smooth muscle subtypes provides insight into embryological origin-dependent disease susceptibility. *Nature Biotechnology* 2012;**30**:165–173.
3. He J, Weng Z, Wu SCM, Boheler KR. Generation of Induced Pluripotent Stem Cells from Patients with COL3A1 Mutations and Differentiation to Smooth Muscle Cells for ECM-Surfaceome Analyses. In: Boheler K, Gundry R, eds. The Surfaceome. New York, NY: Humana Press, 2018:261–302.
4. Yang LB, Gao L, Nickel T, Yang J, Zhou JY, Gilbertsen A, Geng ZH, Johnson C, Young B, Henke C, Gourley GR, Zhang JY. Lactate Promotes Synthetic Phenotype in Vascular Smooth Muscle Cells. *Circ Res* 2017;**121**:1251–1262.
5. Hawthorne RN, Blazeski A, Lowenthal J, Kannan S, Teuben R, DiSilvestre D, Morrissette-McAlmon J, Saffitz JE, Boheler KR, James CA, Chelko SP, Tomaselli G, Tung L. Altered Electrical, Biomolecular, and Immunologic Phenotypes in a Novel Patient-Derived Stem Cell Model of Desmoglein-2 Mutant ARVC. *J Clin Med* 2021;**10**.
6. Yamanaka S, Zahanich I, Wersto RP, Boheler KR. Enhanced Proliferation of Monolayer Cultures of Embryonic Stem (ES) Cell-Derived Cardiomyocytes Following Acute Loss of Retinoblastoma. *PLoS One* 2008;**3**.
7. Zhan M, Riordon DR, Yan B, Tarasova YS, Bruweleit S, Tarasov KV, Li RA, Wersto RP, Boheler KR. The B-MYB Transcriptional Network Guides Cell Cycle Progression and Fate Decisions to Sustain Self-Renewal and the Identity of Pluripotent Stem Cells. *PLoS One* 2012;**7**.
8. Morrissette-McAlmon J, Xu WR, Teuben R, Boheler KR, Tung LS. Adipocyte-mediated electrophysiological remodeling of human stem cell - derived cardiomyocytes. *J Mol Cell Cardiol* 2024;**189**:52–65.
9. Tan JL, Tien J, Pirone DM, Gray DS, Bhadriraju K, Chen CS. Cells lying on a bed of microneedles: an approach to isolate mechanical force. *Proc Natl Acad Sci U S A* 2003;**100**:1484–1489.
10. Fu JP, Wang YK, Yang MT, Desai RA, Yu XA, Liu ZJ, Chen CS. Mechanical regulation of cell function with geometrically modulated elastomeric substrates. *Nat Methods* 2010;**7**:733–736.
11. Shi Y, Sivarajan S, Crocker JC, Reich DH. Measuring cytoskeletal mechanical fluctuations and rheology with active micropost arrays. *Current Protocols* 2022;**2**:e433.
12. Legant WR, Pathak A, Yang MT, Deshpande VS, McMeeking RM, Chen CS. Microfabricated tissue gauges to measure and manipulate forces from 3D microtissues. *Proc Natl Acad Sci U S A* 2009;**106**:10097–10102.
13. Bose P, Huang CY, Eyckmans J, Chen CS, Reich DH. Fabrication and mechanical properties measurements of 3D microtissues for the study of cell-matrix interactions *Methods Mol Biol* 2018;**1722**:303–328.
14. Zhao R, Boudou T, Wang WG, Chen CS, Reich DH. Decoupling cell and matrix mechanics in engineered microtissues using magnetically actuated microcantilevers. *Adv Mater* 2013;**25**:1699–1705.

15. Hawthorne RN, Blazeski A, Lowenthal J, Kannan S, Teuben R, DiSilvestre D, Morrissette-McAlmon J, Saffitz JE, Boheler KR, James CA, Chelko SP, Tomaselli G, Tung L. Altered Electrical, Biomolecular, and Immunologic Phenotypes in a Novel Patient-Derived Stem Cell Model of Desmoglein-2 Mutant ARVC. *J Clin Med* 2021;**10**:3061.
16. Kumar A, D'Souza SS, Moskvin OV, Toh H, Wang BW, Zhang J, Swanson S, Guo LW, Thomson JA, Slukvin II. Specification and Diversification of Pericytes and Smooth Muscle Cells from Mesenchymoangioblasts. *Cell Rep* 2017;**19**:1902–1916.
17. Rzucidlo EM, Martin KA, Powell RJ. Regulation of vascular smooth muscle cell differentiation. *J Vasc Surg* 2007;**45**:25a–32a.
18. Cheung C, Bernardo AS, Pedersen RA, Sinha S. Directed differentiation of embryonic origin-specific vascular smooth muscle subtypes from human pluripotent stem cells. *Nat Protoc* 2014;**9**:929–938.
19. Kilpinen H, Goncalves A, Leha A, Afzal V, Alasoo K, Ashford S, Bala S, Bensaddek D, Casale FP, Culley OJ, Danecek P, Faulconbridge A, Harrison PW, Kathuria A, McCarthy D, McCarthy SA, Meleckyte R, Memari Y, Moens N, Soares F, Mann A, Streeter I, Agu CA, Alderton A, Nelson R, Harper S, Patel M, White A, Patel SR, Clarke L, Halai R, Kirton CM, Kolb-Kokocinski A, Beales P, Birney E, Danovi D, Lamond AI, Ouwehand WH, Vallier L, Watt FM, Durbin R, Stegle O, Gaffney DJ. Common genetic variation drives molecular heterogeneity in human iPSCs *Nature* 2017;**546**:370–375.
20. Topa H, Benoit-Pilven C, Tukiainen T, Pietilainen O. X-chromosome inactivation in human iPSCs provides insight into X-regulated gene expression in autosomes. *Genome Biol* 2024;**25**:144.
21. Raposo AC, Caldas P, Jeremias J, Arez M, Cazaux Mateus F, Barbosa P, Sousa-Luis R, Agua F, Oxley D, Mupo A, Eckersley-Maslin M, Casanova M, Grosso AR, da Rocha ST. Gene reactivation upon erosion of X chromosome inactivation in female hiPSCs is predictable yet variable and persists through differentiation. *Stem Cell Reports* 2025;**20**:102472.
22. MacFarlane EG, Parker SJ, Shin JY, Kang BE, Ziegler SG, Creamer TJ, Bagirzadeh R, Bedja D, Chen YC, Calderon JF, Weissler K, Frischmeyer-Guerrerio PA, Lindsay ME, Habashi JP, Dietz HC. Lineage-specific events underlie aortic root aneurysm pathogenesis in Loeys-Dietz syndrome. *J Clin Invest* 2019;**129**:659–675.
23. Hu YW, Cai ZH, He B. Smooth Muscle Heterogeneity and Plasticity in Health and Aortic Aneurysmal Disease. *Int J Mol Sci* 2023;**24**:11701.
24. Cao G, Xuan X, Hu J, Zhang R, Jin H, Dong H. How vascular smooth muscle cell phenotype switching contributes to vascular disease. *Cell Commun Signal* 2022;**20**:180.
25. Zhang J, McIntosh BE, Wang BW, Brown ME, Probasco MD, Webster S, Duffin B, Zhou Y, Guo LW, Burlingham WJ, Kent C, Ferris M, Thomson JA. A Human Pluripotent Stem Cell-Based Screen for Smooth Muscle Cell Differentiation and Maturation Identifies Inhibitors of Intimal Hyperplasia. *Stem Cell Reports* 2019;**12**:1269–1281.

### SUPPLEMENTAL FIGURES AND FIGURE CAPTIONS

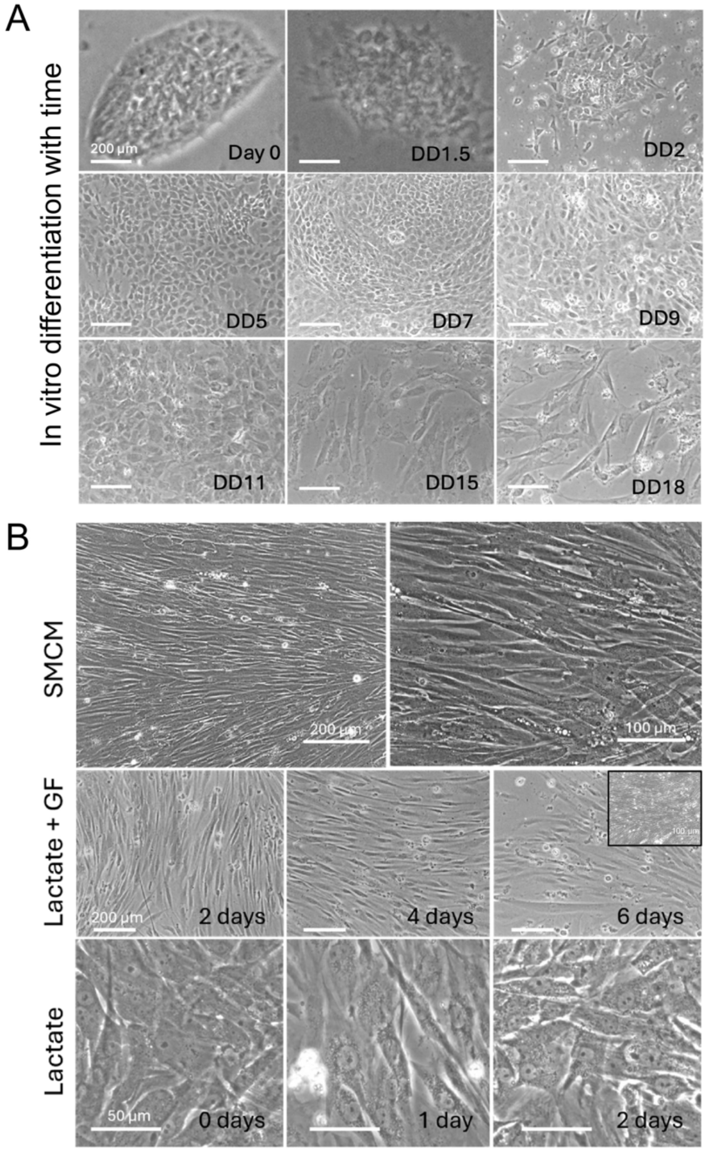

**Figure S1.** Morphological characteristics of in vitro differentiating and vSMC plated under different conditions. A) Representative images of hPSC line WTC11 differentiated into lineage-specific (paraxial mesoderm) vSMC at selected time points as indicated in the figure. At differentiation Day (DD) 18, cells were transitioned and cultivated in SMCM until confluent, at which time the cells were passaged (p1). Scale bar 200 µm. B**)** Representative images of in vitro derived vSMC enriched with lactate-containing media vs SMCM. Top: Brightfield images of JHU001-derived vSMC cultivated to confluency in SMCM. Scale bar 200 and 100 µm. Middle: Brightfield images of cells cultivated to confluency and then incubated with 4 mM lactate + GF medium for 6 days. The inset at day 6 shows a culture with high density after selection. Scale bar 200 µm. Bottom: Brightfield images of vSMC cultivated to confluency and then incubated for 48 hours in 4 mM lactate medium (Lac) that did not contain any GFs. Scale bar 50 µm.

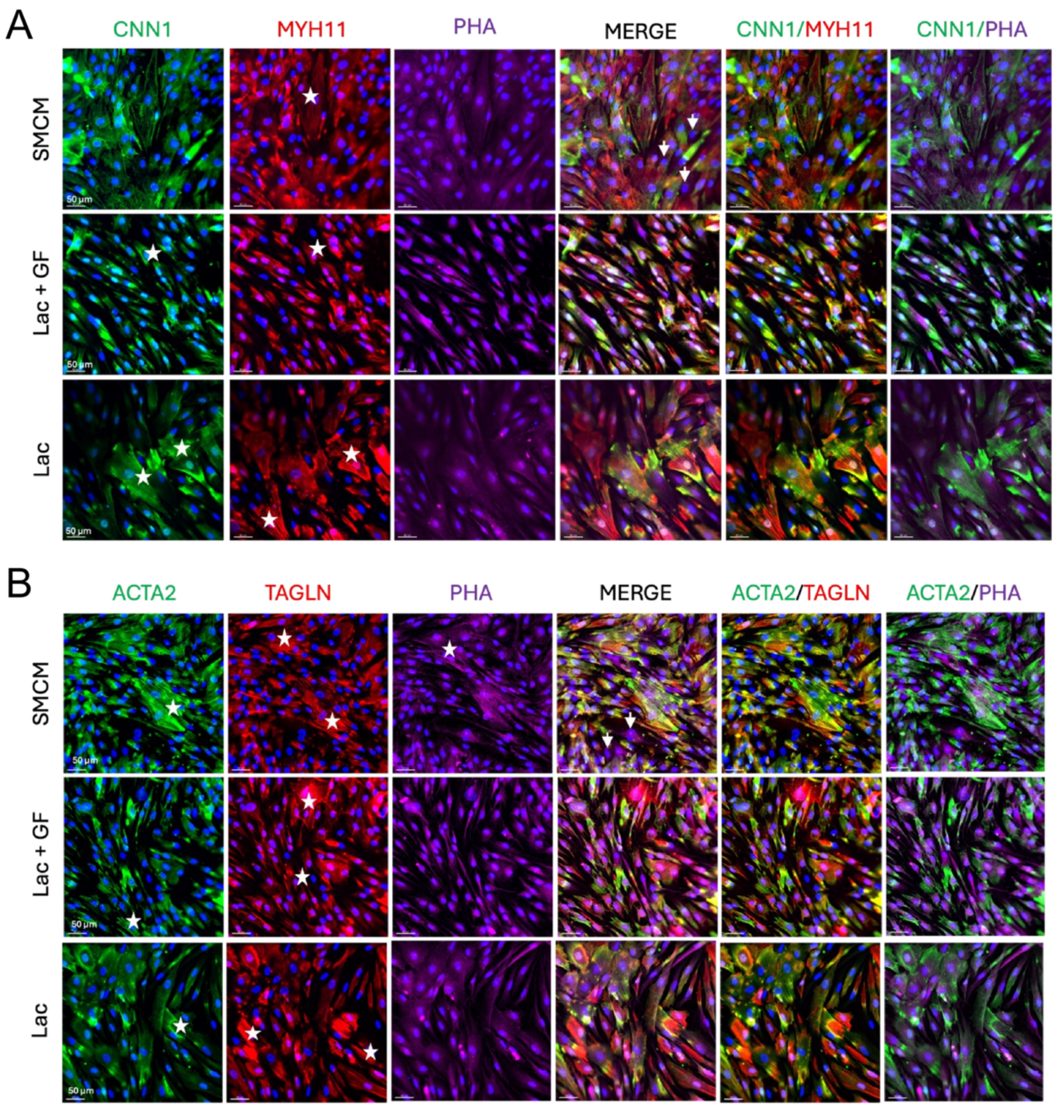

**Figure S2.** Representative immunofluorescent images of JHU001 hPSC-derived vSMC cultivated in SMCM or enriched with lactate media. A) Immunofluorescent (IF) images and merged images show IF staining of proteins with antibodies to CNN1, MYH11, PHA and the nuclear dye DAPI. Note, the MYH11 specific antibody used on in fixed cells likely shows some cross-reactivity with other myosin proteins or some non-specificity (see main text, section 3.2). Cells were cultivated in SMCM to near confluency followed by 3 additional days of cultivation in SMCM or were grown to near confluency and enriched by cultivation for 6 days in a 4 mM lactate medium containing growth factors (Lac + GF) and one additional day in SMCM or were grown to near confluency and enriched for 2 days a 4 mM lactate medium lacking growth factors (Lac) and one additional day in SMCM. White arrows in the merged image indicate cells positive for PHA but lacking CNN1 or MYH11. B) IF images and merged images of hiPSC-derived vSMC stained for proteins with antibodies to ACTA2, TAGLN, PHA, and with nuclear dye DAPI. White arrows in the merged image indicate cells positive for PHA but lacking ACTA2 or TAGLN. Stars highlight cells with fiber-like structures in S3A and S3B. Scale bar 50 μm.

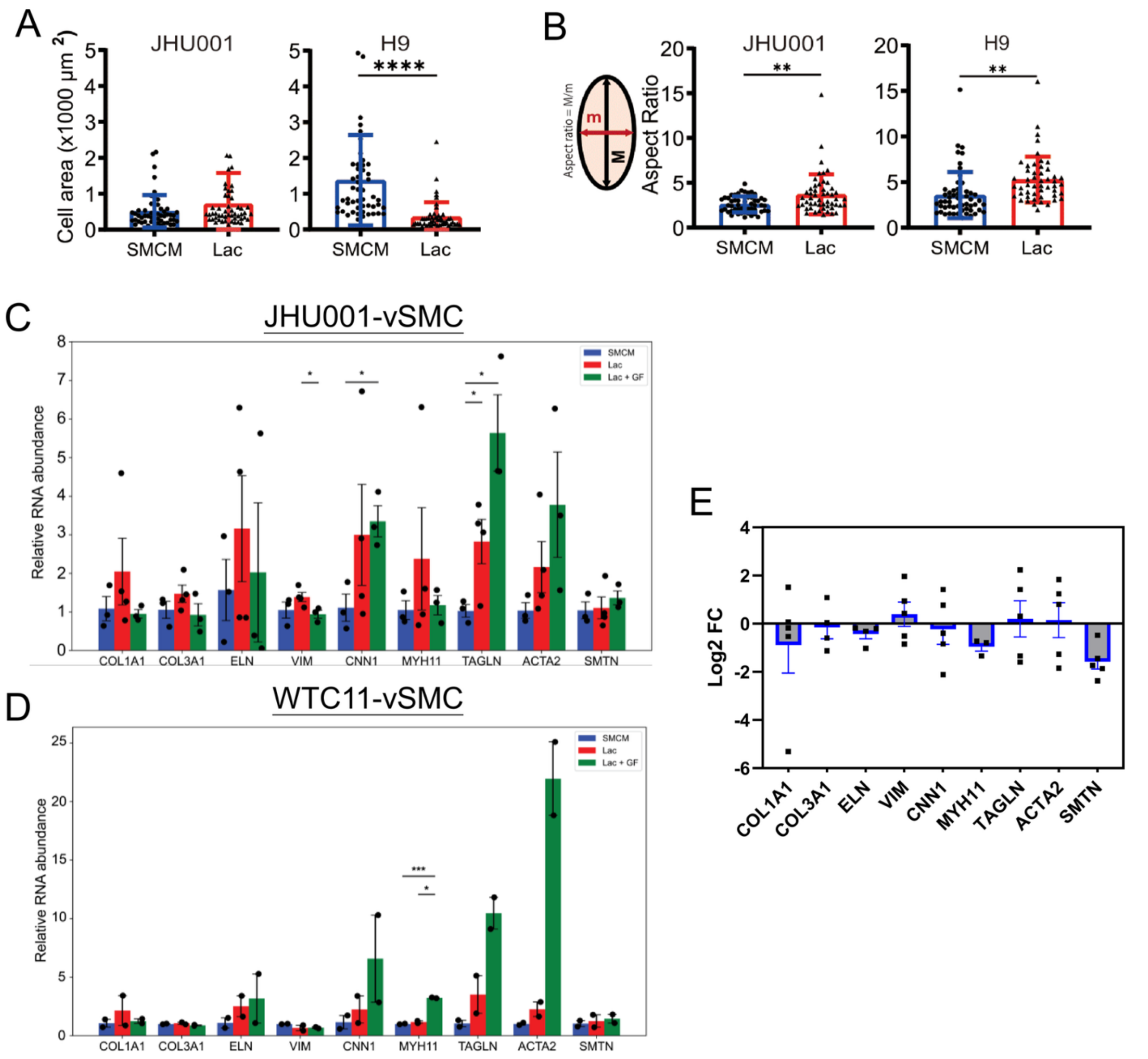

**Figure S3.** Characteristics of in vitro differentiated, PM-derived hPSC-vSMC. A) Assessment of cell area B) elongation (aspect ratio of best-fit ellipse) of JHU001- and H9-derived vSMC treated with Lac medium vs SMCM. Error bars: ±SD. JHU001: SMCM n=53, Lac n=51; H9 SMCM n=50; Lac n=50. Relative transcript abundance of ECM, cytoskeletal and contractile proteins in hPSC-vSMC cultivated in SMCM, Lac or Lac + GF. C) qPCR analyses of transcripts from JHU001 hPSC-vSMC (n=3-4). D) qPCR analyses of transcripts from WTC11 hPSC-vSMC (n=2). Error bars for C and D: ±SEM. E) qPCR data generated using TaqMan probes with H9-vSMC. Data are presented as $\log_2$ fold change (Lac vs. SMCM) and shown as mean ± SEM (n=3-5). Standard Abbreviations: COL1A1 – collagen type I, alpha 1; COL3A1 – collagen type III, alpha 1; ELN – elastin; VIM – vimentin; CNN1 – calponin 1; MYH11 – smooth muscle myosin heavy chain 11; TAGLN – transgelin; ACTA2 – smooth muscle actin; SMTN – smoothelin. Statistical analysis was carried out using a Welch's *t*-test in A and B. $P<0.05$ was considered statistically significant (*p<0.05, **p<0.01, ***p<0.001).

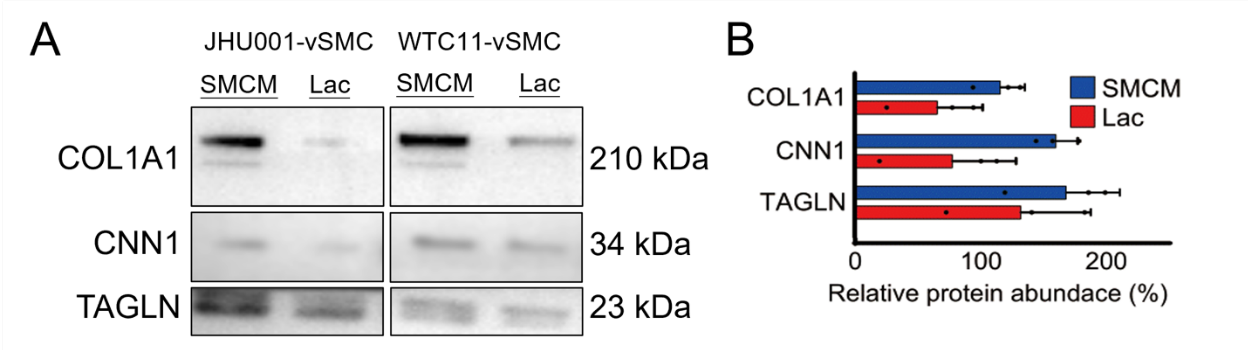

**Figure S4**. Representative Western blot results of JHU001- and WTC11-derived vSMC under SMCM and Lac conditions. (A) Composite images of immunoblots showing COL1A1, CNN1 and TAGLN protein levels in vSMC. Molecular weights (kDa) for the corresponding protein are shown on the right of each gel based on size markers. (B) Bar graph showing the quantification of COL1A1, CNN1 and TAGLN protein levels in samples cultured in SMCM or Lac medium and following normalization for protein loading from all samples (JHU001 and WTC11). Data are presented as a percentage (%). Blue bars, SMCM; red bars, Lac. Error bars: ± SEM, n=3.

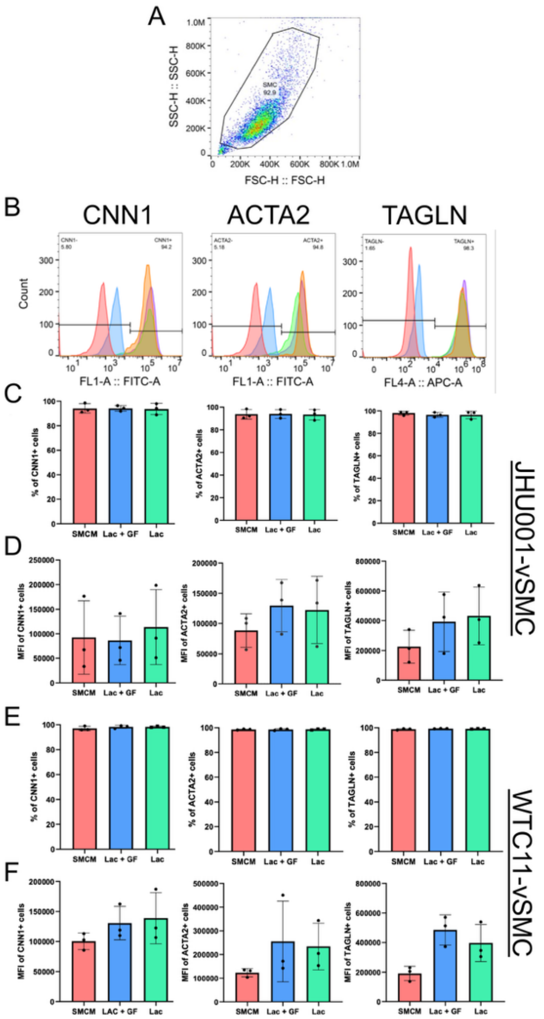

**Figure S5.** Flow cytometry analysis of selected contractile proteins in hiPSC-vSMC cultured in SMCM or lactate medium. A) Representative gating of analyzed cell populations for fixed JHU001 vSMC plotted as forward scatter (FSC) vs side scatter (SCC). B) Histograms showing the signal distributions of unstained cells (blue) and isotype control antibody (red) labelling across a population of cells. Each histogram also shows the fluorescence intensity of vSMC positive markers for CNN1, ACTA2 or TAGLN in cells cultured with SMCM (green), Lac + GF (orange), and Lac medium (violet). C) Bar graphs showing the percentages of gated JHU001-vSMC with positive signals for the indicated protein, based on the gating parameters described in B) and performed on independent batches of in vitro differentiated vSMC. D) The mean fluorescence intensity (in arbitrary units) of the fluorescent signals generated in the selected population of JHU001-vSMC. E) Graphical representation of the percentages of gated WTC11-vSMC with positive signals for the indicated protein. F) The mean fluorescence intensity (in arbitrary units) of the fluorescent signals generated in the selected population of WTC11-vSMC. An increase in MFI was considered indicative of an increase in total marker protein in the population of cells. Error bars ± SD (n=3). Statistical analysis was carried out using a one-way ANOVA in C, D, E and F. $P$<0.05 was considered statistically significant, but in these experiments, no significant difference could be demonstrated.

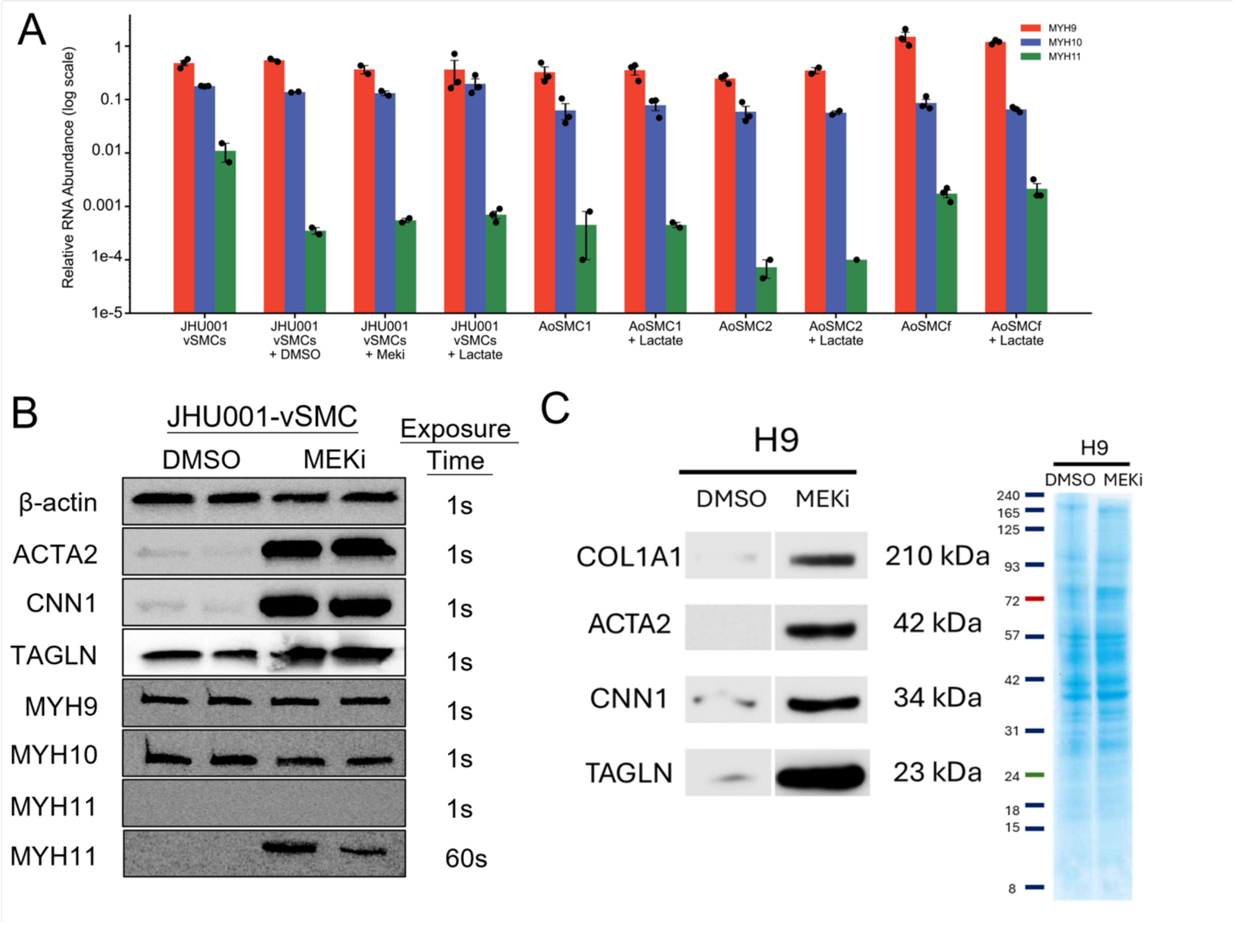

**Figure S6.** Assessment of non-muscle and smooth muscle myosin heavy chain transcripts and proteins, and Data from H9-derived vSMC. A) qPCR analysis of non-muscle (MYH9, MYH10) and smooth muscle myosin heavy (MYH11) in JHU001 hPSC-vSMC and AoSMC (as indicated). Data are presented as $2^\Delta\Delta Ct$ on a log scale after normalization to RPL32, n=2-3. B) Western blot composite showing the presence of non-muscle and smooth muscle myosin heavy chains in JHU001-vSMC from two independent batches of treated cells. In DMSO-treated control vSMC, MYH11 was undetectable, while MYH9 and MYH10 were abundant. In MEKi-treated, contractile vSMC, all three myosin heavy chain isoforms were detectable; however, MYH11 could only be detected after relatively long exposure times. C) Representative Western blot composite images and Coomassie blue staining of whole membrane following protein transfer of lysates prepared from DMSO and MEKi-treated H9-derived vSMC.

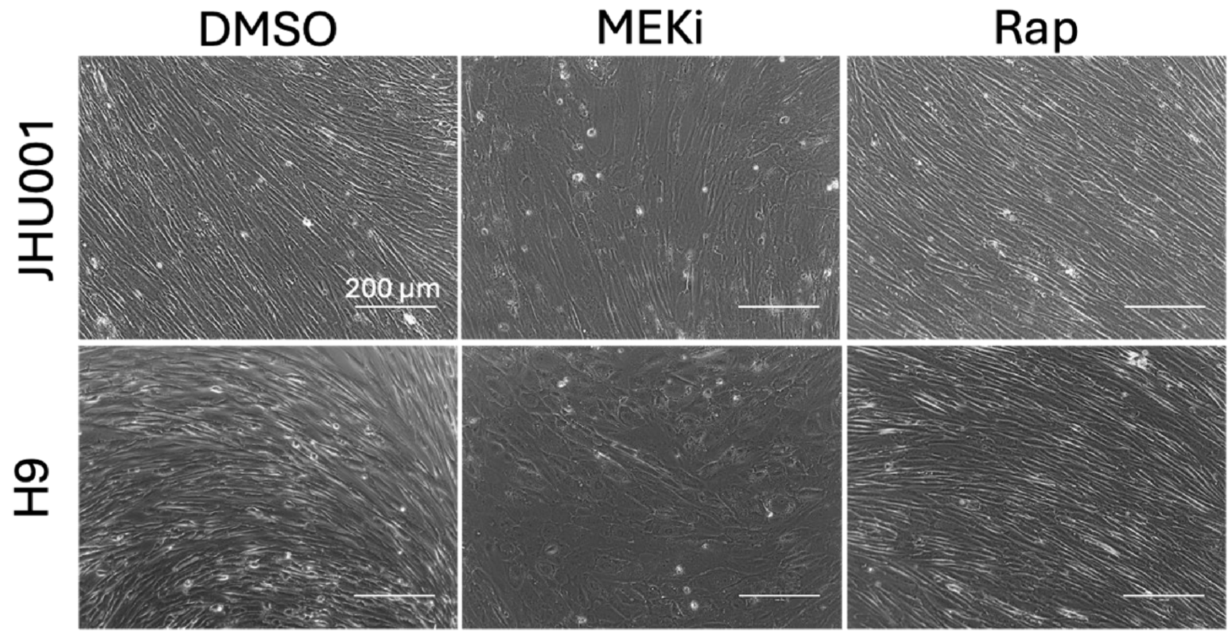

**Figure S7.** Representative hPSC-vSMC images of cells treated with DMSO, MEKi, and Rap. Brightfield images of JHU001 and H9-derived vSMC imaged after 8 days of treatment with DMSO, 1 µM MEKi, and 5 nM Rap. Scale bar 200 mm. Cells treated with DMSO and Rap were spindle-shaped and generally overgrown following extended treatment times.

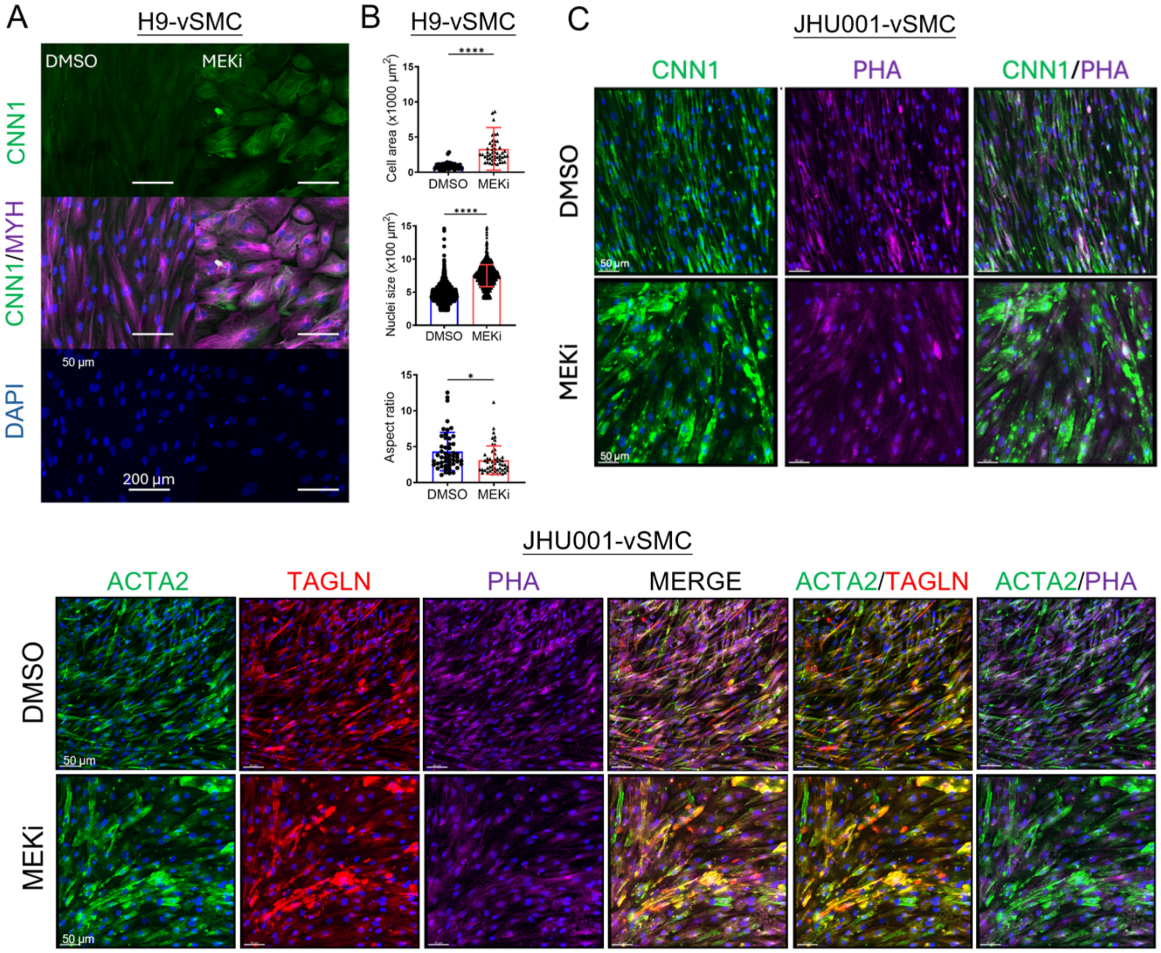

**Figure S8**. Phenotypic traits of DMSO vs MEKi-treated vSMC from hPSC lines. A) Immunofluorescence staining and physical characteristics of DMSO- and MEKi-treated vSMC from H9 human embryonic stem cells. IF images of cells following 1 µM MEKi treatments for 6 days. Cells were passaged onto coverslips, allowed to attach for ~18 hours, fixed and stained with antibodies to CNN1 and MYH11 (shown as non-specific MYH – see main text, section 3.2). Scale bar 200 µm. B) Physical characteristics of control and MEKi-treated H9-derived vSMC, as indicated in the bar graphs. Data are shown as mean±SD, n = 50. C) Immunofluorescence images of JHU001-derived vSMC-restricted contractile proteins (CNN1, ACTA2, F-actin, TAGLN) in monolayer cultures after 6 days of treatments (DMSO or MEKi) and 2 days in SMCM. The staining intensities show contractile protein signal heterogeneity as a function of treatment. The merged panels and pairwise overlaps illustrate the signal heterogeneity among the different proteins. Scale bar 50 µm. Statistical analysis was carried out using a Student's *t*-test in B. $P<0.05$ was considered statistically significant (*$p<0.05$, ****$p<0.0001$).

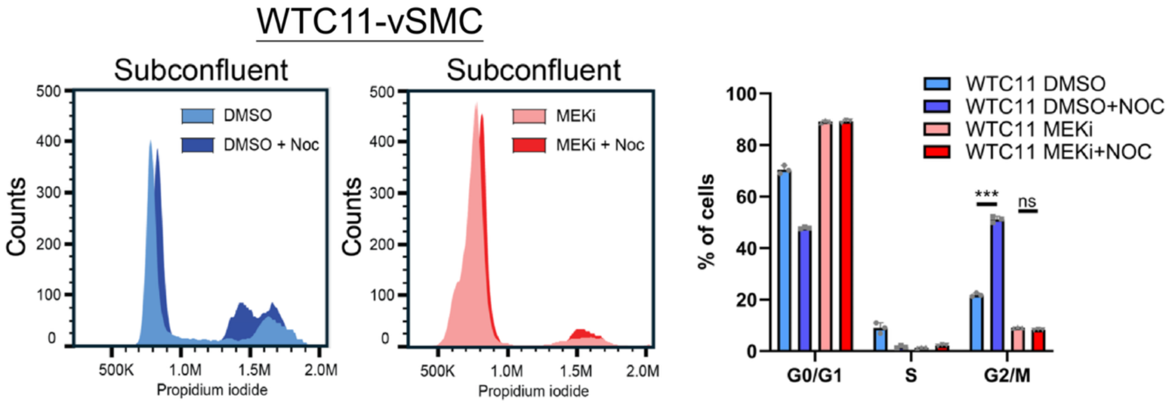

**Figure S9.** Cell cycle analysis of WTC11-vSMC. Subconfluent vSMC were examined after cultivation in DMSO and MEKi ± 10 µM Nocodazole (Noc). The bar graphs show that cells grown in SMCM represent a generally proliferating, synthetic vSMC phenotype, while those treated with MEKi are more consistent with a non-proliferating, contractile vSMC phenotype. Bars represent mean ±SD (n=3). Statistical analysis was carried out using a two-way ANOVA. *P*<0.05 was considered statistically significant (***p<0.001).

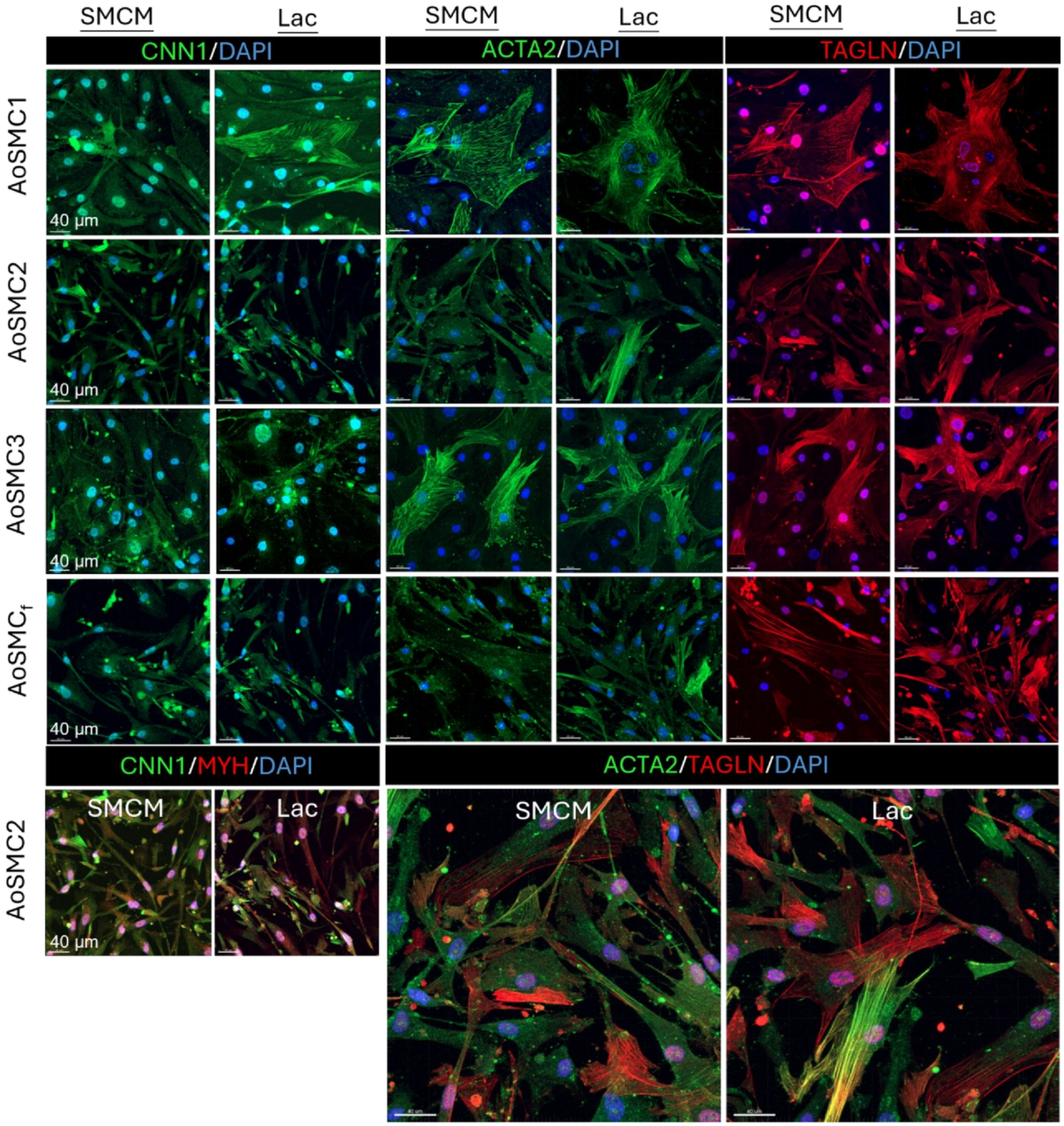

**Figure S10.** Representative IF images of commercially available vSMC derived from human aorta. Immunofluorescent images are shown for near confluent cultures of 3 batches of AoSMC and 1 batch of fetal AoSMC in SMCM following 48 hours of selection in Lac Medium. Merged images showing individual protein signal heterogeneities within cells for CNN1, MYH, ACTA2 and TAGLN (see the lower panels). Scale bar 40 μm.

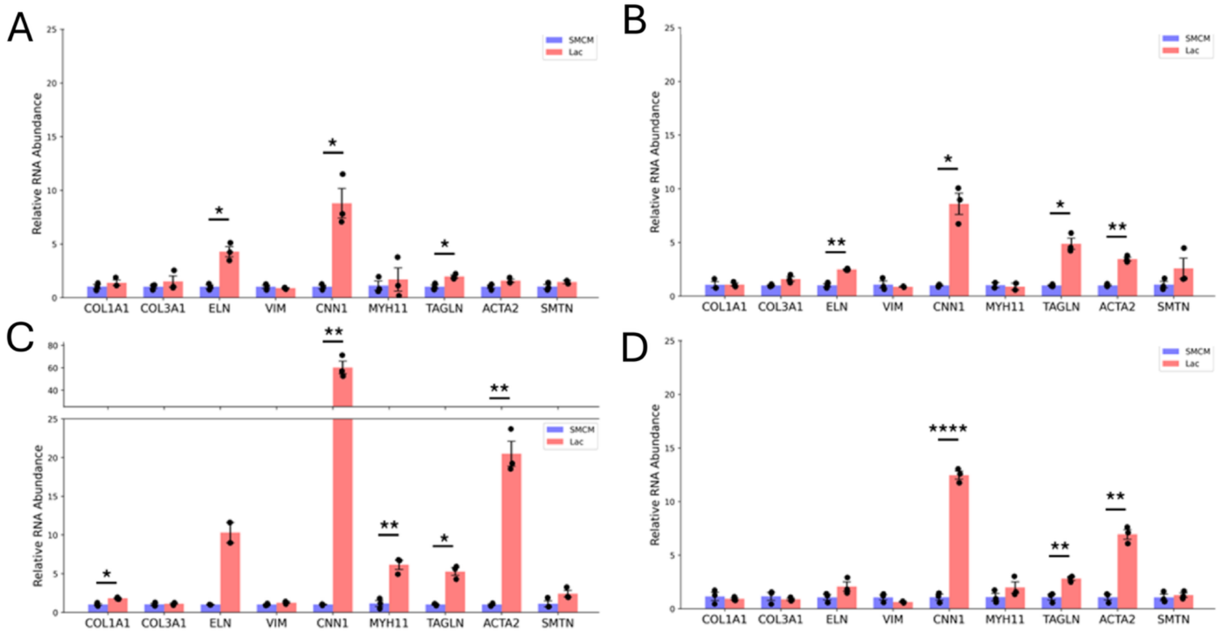

**Figure S11.** Relative RNA abundance in primary AoSMC cultivated in Lac medium compared to SMCM. A) Transcript data for selected transcripts in AoSMC1. B) Transcripts expressed in line AoSMC2. C) Transcripts expressed in line AoSMC3. D) Transcripts expressed in fetal line AoSMC$_f$. RNA abbreviations are defined in Figure S4. Statistical analysis was carried out using a Welch's *t*-test in A, B, C and D. Error bars: ± SEM, n=2-3. *P*<0.05 was considered statistically significant (*p<0.05, **p<0.01, ***p<0.001).

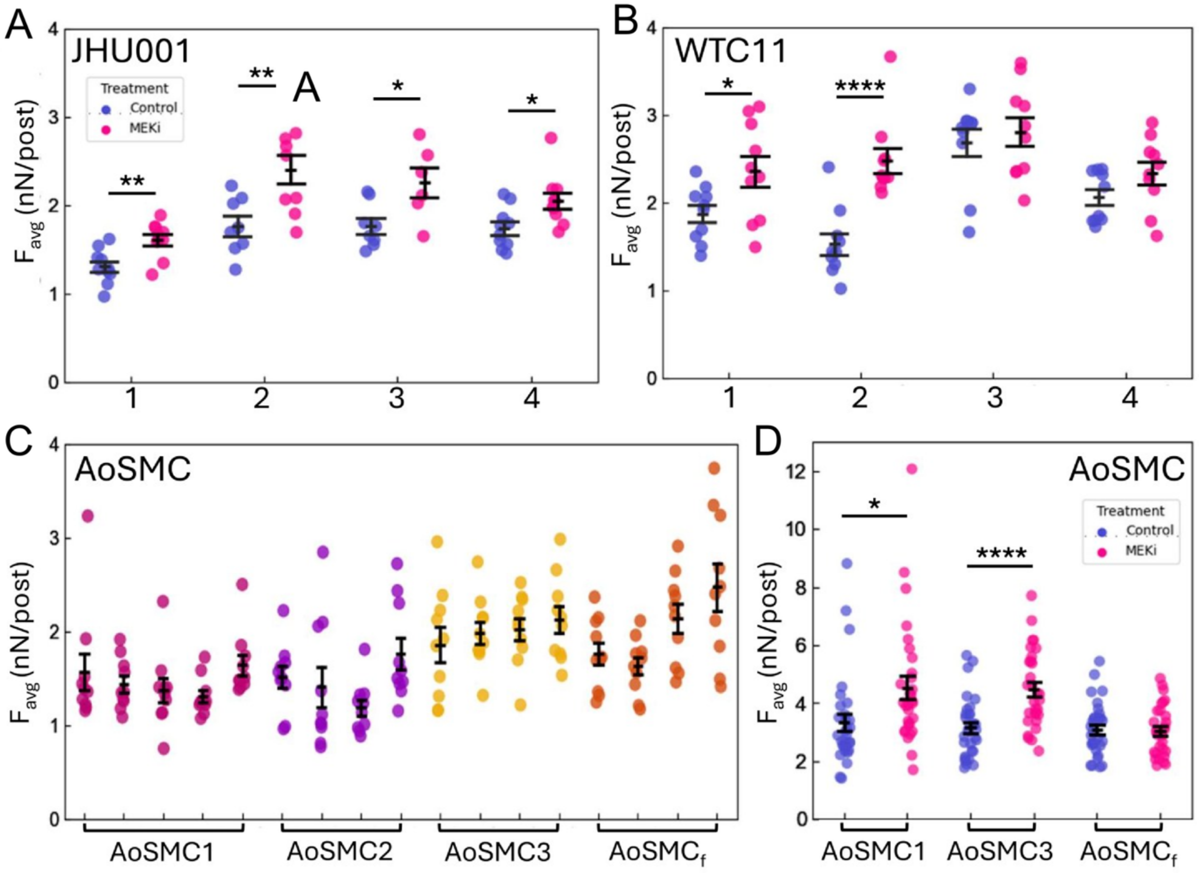

**Figure S12.** Assessment of hPSC-derived (±MEKi treatment) and primary AoSMC single-cell contractility. A) Average force per post for JHU001-derived vSMC ± MEKi treatment for cells. B) Average force per post for WTC11-derived vSMC ± MEKi treatment. All experiments with hPSC-vSMC were performed on cells from four independent differentiations. C) Average force per post measured for primary AoSMC from four different commercially available lines, including one fetal line. At least four independent experiments were conducted with each line, where each experiment was performed on cells from different frozen stocks or passage numbers. Measurements were made using mPADs to assess single-cell contractile forces. Measurements across different experiments show significant variability, even within the same cell line. D) Effects of MEKi treatment on AoSMC contractility. Error bars: ±SEM, n=6-11. Statistical analysis was carried out using a Welch's *t*-test in A and B. $P<0.05$ was considered statistically significant (*p<0.05, **p<0.01, ****p < 0.0001). The coefficient of variation (CV) across the independent experiments with each cell line in Panels A-C above are, with CV = σ/μ: Control JHU001-vSMC (N = 4) $CV_C$ = 12.2%; MEKi-treated JHU001-vSMC (N = 4): $CV_M$ = 15.8%; $AoSMC_1$ (N = 5): $CV_1$ = 8.4%; $AoSMC_2$ (N = 4): $CV_2$ = 14.0%; $AoSMC_3$ (N = 4): $CV_3$ = 4.7%; $AoSMC_f$ (N = 4): $CV_4$ = 16.4%.

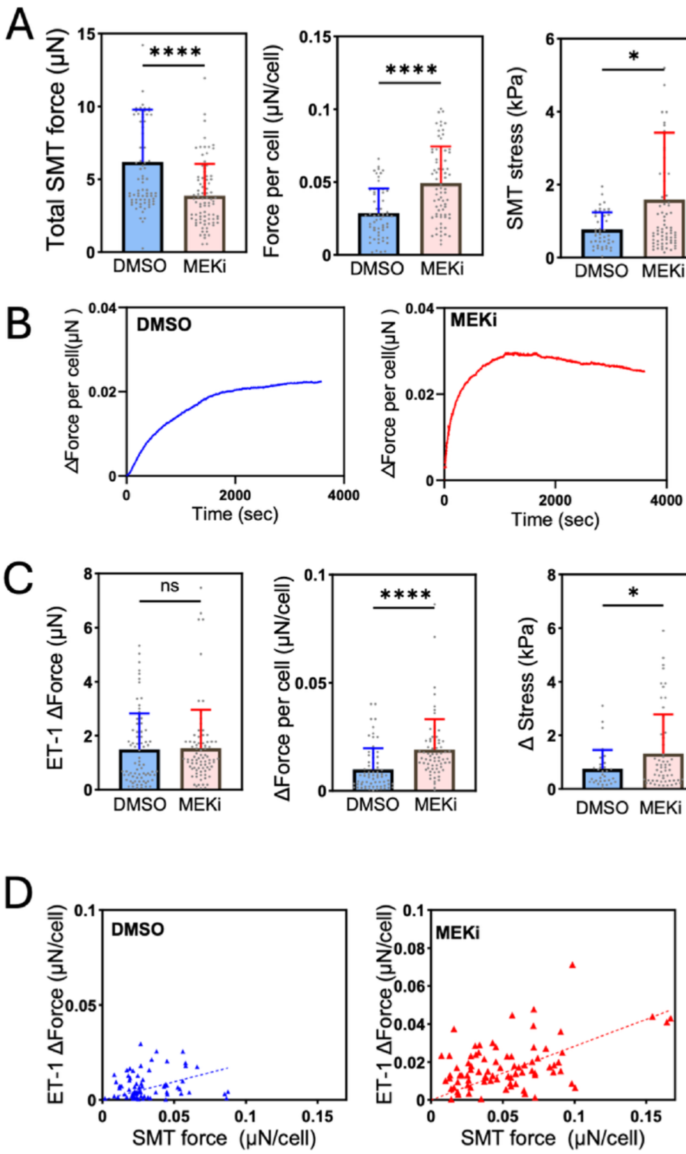

**Figure S13.** Contractile force analysis of WTC11-vSMC microtissues treated with DMSO or MEKi for 4 days obtained using μTUG arrays with pillar spring constant k = 0.25 μN/μm. (A) Basal SMT force was 3.12 ± 1.62 μN (n = 68) in DMSO vs. 3.72 ± 2.11 μN (n = 93) in MEKi. Force per cell was 0.024 ± 0.012 μN (n = 53) vs. 0.042 ± 0.030 μN (n = 100). Stress was 0.45 ± 0.26 μN/μm² (n = 68) vs. 1.23 ± 0.73 μN/μm² (n = 91). (B) Representative traces of SMT force generation following exposure to 1 μM ET-1. (C) ET-1–induced responses after 1 h with 10 μM ET-1 in DMSO vs. MEKi: Δ force was 2.00 ± 1.05 μN (n = 68) vs. 2.21 ± 1.18 μN (n = 88); Δ force per cell was 0.018 ± 0.010 μN (n = 73) vs. 0.034 ± 0.017 μN (n = 100); Δ stress was 0.29 ± 0.19 μN/μm² (N = 62) vs. 0.85 ± 0.47 μN/μm² (n = 90). Western blot analysis of selected ECM and contractile proteins in WTC11-vSMC ± MEKi. (D) Correlation between baseline SMT force and ET-1–induced Δ force, showing similar regression slopes in DMSO (n = 46, slope = 0.18, $R^2$ = 0.14) and MEKi (n = 72, slope = 0.24, $R^2$ = 0.05.) All data are shown as mean ± SD. Statistical analysis was carried out using an independent two-sample Student's *t*-test in A and C. *P*<0.05 was considered statistically significant (*p<0.05, ****p<0.0001).

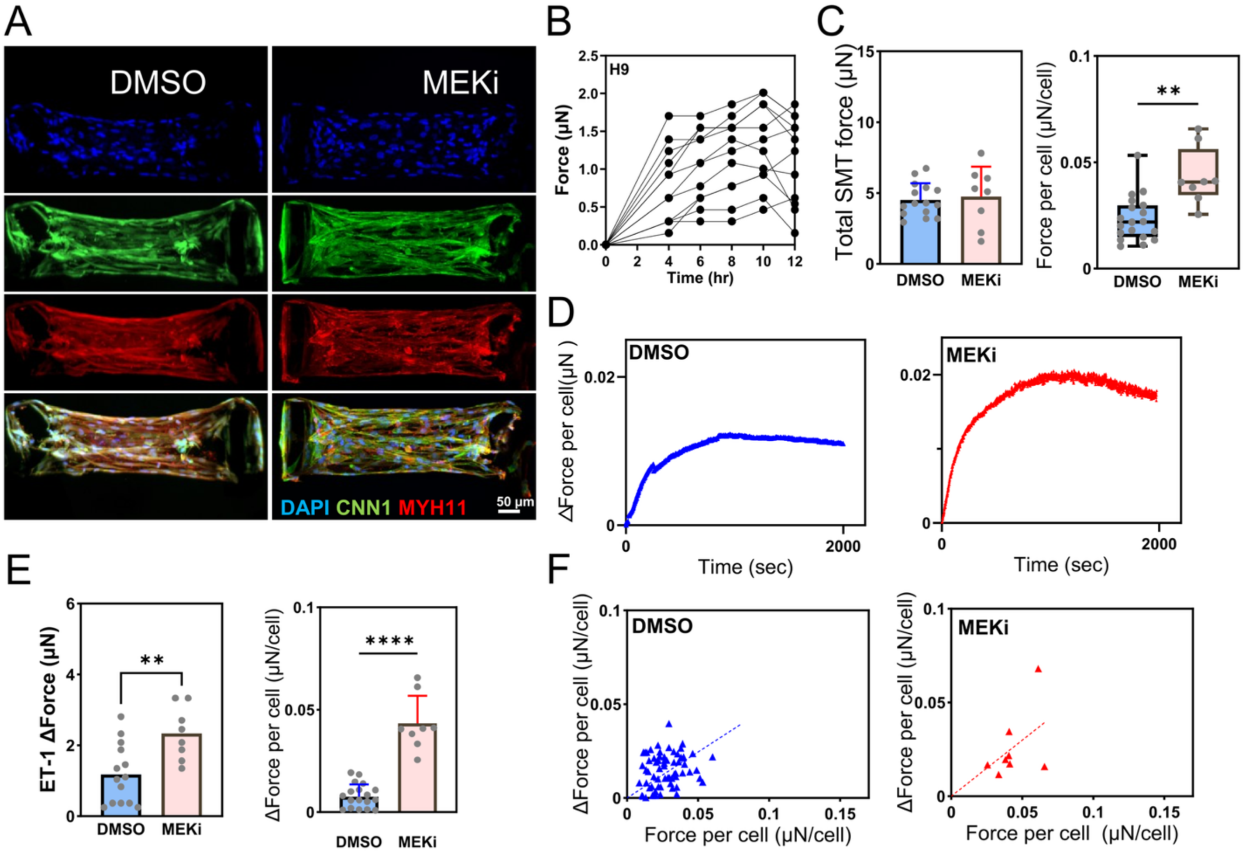

**Figure S14.** Force measurements of humanized microtissues composed of H9-derived vSMC. A) Immunostaining showing the presence of CNN1 and non-specific MYH11 aligned along the long axis of the SMTs. Nuclei are stained with DAPI. Scale bar 50 µm. B) Representative traces of microtissue force vs time of compaction. C) Graphs of microtissue forces for SMTs treated with DMSO and MEKi and obtained from µTUGs (k = 0.25 µN/µm). Total SMT force was 4.51 ± 1.17 µN vs. 4.75 ± 2.11 µN (ns), and the force per cell was 0.02 ± 0.01 µN vs. 0.04 ± 0.01 µN ($p$<0.01). Error bars: ± SD; DMSO n=15, MEKi n=8. D) Traces showing maximum force generation in SMTs following application of 1 µM ET-1. E) Maximum force generation in vSMC ± ET-1. Forces were 1.18 ± 0.84 µN in DMSO vs. 2.34 ± 0.75 µN in MEKi (p<0.01); Δ force per cell was 0.008 ± 0.001 µN vs. 0.04 ± 0.004 µN ($p$<0.0001); Error bars: ±SD; DMSO n=19, MEKi n=8. F) Scatter plot showing the correlation between ET-1 response and baseline SMT force. DMSO: slope = 0.49, $R^2$=0.14, n=72. MEKi: slope = 0.59, $R^2$=0.24, n=8. Statistical analysis was carried out using a Student's t-test in C, E and F. P<0.05 was considered statistically significant (**p<0.01, ****p<0.0001).

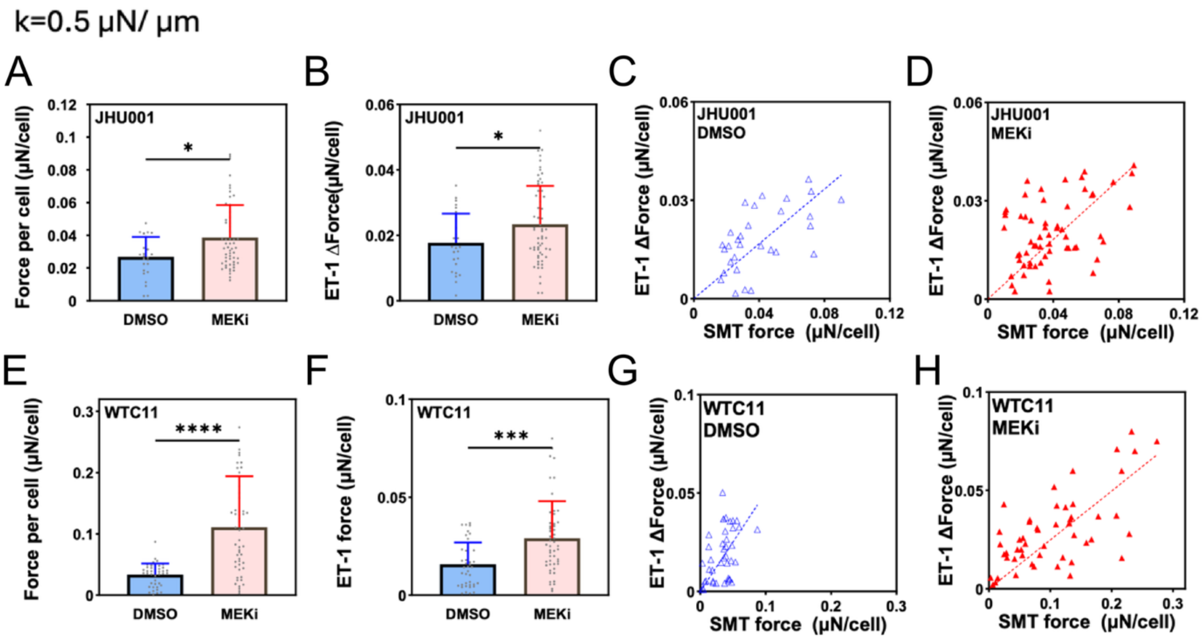

**Figure S15.** Developed force and response to ET-1 for JHU001 and WTC11 SMTs on uTUGs with pillar spring constants of k = 0.5 μN/μm. (A) JHU001 baseline force per cell: DMSO 0.027 ± 0.012 μN/cell (n = 26) vs MEKi 0.038 ± 0.019 μN/cell (n = 25), $p$ = 0.0279; (B) JHU001 ET-1 ΔForce: DMSO 0.018 ± 0.008 μN/cell (n = 26) vs MEKi 0.023 ± 0.01 μN/cell (n = 25), $p$ = 0.049; (C) JHU001 DMSO correlation (ET-1 response vs baseline): $N$ = 26, slope = 0.41, $R^2$ = 0.20; (D) JHU001 MEKi correlation: $n$ = 25, slope = 0.45, $R^2$ = 0.14; (E) WTC11 baseline force per cell: DMSO 0.03 ± 0.018 μN/cell (n = 35) vs MEKi 0.11 ± 0.08 μN/cell (n = 53), $p$< 0.0001 (exact $p$ = 4.5 × $10^{-8}$); (F) WTC11 ET-1 ΔForce: DMSO 0.015 ± 0.01 μN/cell (n = 35) vs MEKi 0.029 ± 0.018 μN/cell (n = 53), $p$ = 0.0001; (G) WTC11 DMSO correlation: $n$= 35, slope = 0.50, $R^2$ = 0.15; (H) WTC11 MEKi correlation: $n$ = 53, slope = 0.20, $R^2$ = 0.34. Data are presented as mean ± SD. Statistical analysis was carried out using an independent two-sample Student's $t$-test for group comparisons (A, B, E, F), and correlation statistics (C, D, G, H) were determined using linear regression. $P$<0.05 was considered statistically significant (*p<0.05, ***p<0.001, ****p<0.0001).

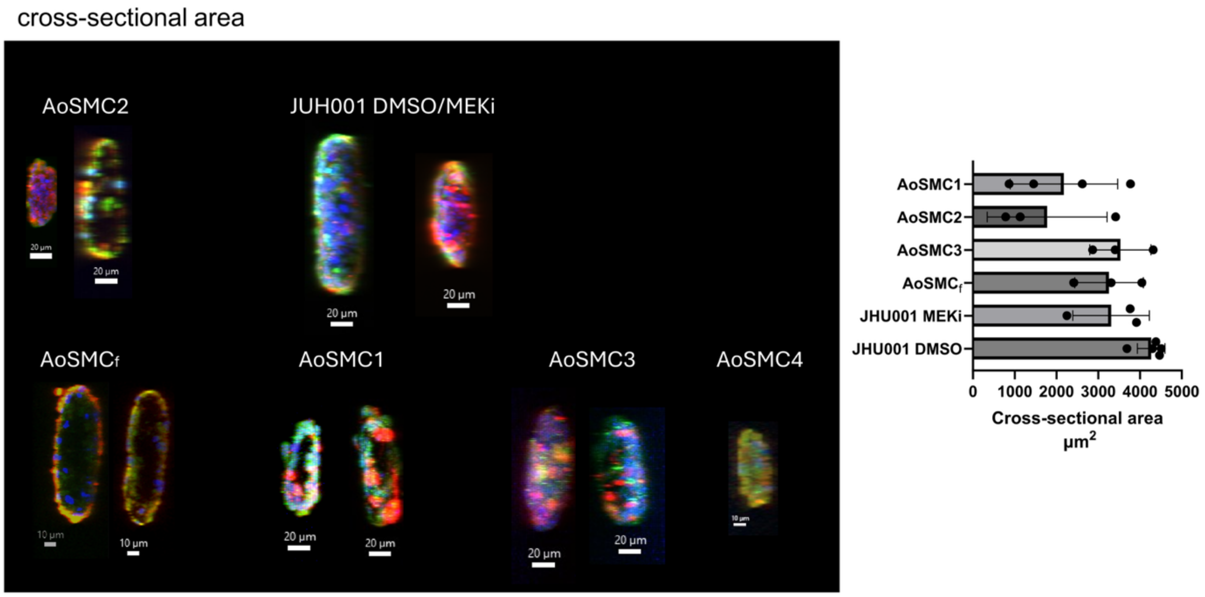

**Figure S16.** Cross-sectional area of smooth muscle microtissues (SMTs) for determining SMT stress. (A) Confocal images showing SMT cross sections. Green: CNN1; Red: MYH; Blue: DAPI (nuclei). Scale bars: 10 or 20 µm, as indicated. A video showing a Z-stack image composite of an AoSMC is shown in Supplemental Video 1. (B) Graphic showing averaged cross-sectional areas of hPSC-derived SMTs and primary AoSMTs. JHU001-SMT DMSO: 4270 ± 330 µm² (n = 5); JHU001-SMT MEKi: 3310 ± 920 µm² (n = 3); $AoSMC_f$: 3260 ± 818 µm² (n = 3); AoSMC3: 3530 ± 740 µm² (n = 3); AoSMC2: 1780 ± 1430 µm² (n = 3); AoSMC1: 2180 ± 1290 µm² (n = 4). Data are shown as mean ± SD with individual values overlaid.

### SUPPLEMENTARY TABLES

**Table S1.** Primers for qPCR analysis of vSMC and SMTs

a. Quantitative PCR (qPCR) primers

| Target | Forward primer | Reverse primer |
|---|---|---|
| **ACTA2**<br>actin, alpha 2, smooth muscle, aorta | GTGTTGCCCCTGAAGAGCAT | GCTGGGACATTGAAAGTCTCA |
| **COL1A1**<br>collagen type I alpha 1 | CCCCGAGGCTCTGAAGGTC | GGA GCA CCA TTG GCA CCT TT |
| **COL3A1**<br>collagen type III alpha 1 chain | GCAGGGTCTCCTGGTTCAAA | CGGGACCCATTTCGCCTTTA |
| **CNN1**<br>calponin 1 | CTGAGAGAGTGGATCGAGGG | CTGGCTGCAGCTTATTGATG |
| **ELN**<br>elastin | TTCCCCCCAGTTACCTTT | CTAACCCACCAACTCCTGGG |
| **MYH9**<br>non-muscle myosin heavy chain 9 | CAGCAAGCTGCCGATAAGTAT | CTTGTCGGAAGGCACCCAT |
| **MYH10**<br>non-muscle myosin heavy chain 10 | GCAGGAGAACACCTAAAGTCTG | TGTCCCGGAATAGGAATATAGCC |
| **MYH11**<br>myosin heavy chain 11 | CGACATGTACAAGGCCAAGGA | TAGAATGGACTGGTCCTCCC |
| **RPL32**<br>ribosomal protein L32 | AGTGCCTAGTATTCTGCCAGC | AGAGTGTCTTCCAATCGCCAG |
| **SMTN**<br>smoothelin | TCCCGCACATTCTTGAGCATT | TGGAACTGGAACTGGGGACC |
| **TAGLN**<br>transgelin | CCGTGCACATCCCAACTGC | CCATCTGAAGGCCAATCACAT |
| **VIM**<br>vimentin | GACGCCATCAACACCGAGTTC | CTTTGTCGTTGGTTACCTGGT |

b. Taqman primer assay information for primers and probes

| **Target** | **Assay ID** |
|---|---|
| ACTA2 | Hs00426835_g1 |
| CNN1 | Hs00154543_m1 |
| MYH11 | Hs00224610_m1 |
| SMTN | Hs00199489_m1 |
| TAGLN | Hs01038777_g1 |
| COL1A1 | Hs00164004_m1 |
| COL3A1 | Hs00943809_m1 |
| GADPH | Hs02758991_g1 |
| VIM | Hs00958111_m1 |
| ELN | Hs00355783_m1 |

**Table S2.** List of antibodies for immunofluorescence (IF), Western blots (WB) and flow cytometry.

| Primary Antibody | Source | Catalog No. | Species | Application | Dilution |
|---|---|---|---|---|---|
| MYH11 | Abcam | 82541 | Rabbit | IF<br>WB | 1:100<br>1:1000<br>1:2000 |
| MYH11 | Abcam | 133567 | Rabbit | Flow | 1:50 |
| MYH10 | Cell Signaling | 49349S | Rabbit | WB | 1:1000 |
| MYH9 | Cell Signaling | 8824S | Rabbit | WB | 1:1000 |
| ACTA2 (α-SMA) | Abcam | 7817 | Mouse IgG2a | IF<br>WB | 1:100<br>1:3000 |
| CNN1 | SIGMA | c2687 | Mouse | IF<br>Flow<br>WB | 1:100<br>1:200<br>1:5000 |
| TAGLN (SM22α) | Abcam | 14106 | Rabbit | IF<br>WB | 1:100<br>1:1000 |
| COL1A1 | Santa Cruz Biotechnology | SC-293182 | Mouse | Flow<br>WB | 1:100<br>1:200 |
| Vimentin | Santa Cruz Biotechnology | SC-6260 | Rabbit | Flow | 0.5 μL/$10^5$ cell |
| Secondary antibody | Source | Catalog No. | Species | Application | Dilution |
| Alexa Fluor 647 goat anti-rabbit IgG (H+L) | Invitrogen | A21245 | Goat | IF<br>Flow | 1:500<br>1:1000 |

| | | | | | |
|---|---|---|---|---|---|
| Alexa Fluor 488 goat anti-mouse IgG (γ1) | Invitrogen | A21121 | Goat | IF<br>Flow | 1:500<br>1:1000 |
| F(ab')2-Goat anti-Mouse IgG (H+L) Cross-Adsorbed Secondary Antibody, Alexa Fluor Plus 488 | Invitrogen | A48286TR | Goat | IF | 1:500 |
| Alexa Fluor™ 488 goat anti-mouse IgG3 | Invitrogen | A24877 | Mouse IgG3 | Flow | 0.2 µL/50 µL |
| Goat anti-rabbit IgG HRP | GeneTex | GTX213110-01 | Goat IgG | WB | 1:5000 |
| Goat anti-mouse IgG HRP | SeraCare | 5220-0460 | Goat IgG (γ) | WB | 1:10000 |
| Isotype control and DNA Dye | Source | Catalog No. | Species | Application | Dilution |
| Mouse isotype Control | Bio-RAD | MCA928 | Mouse IgG1 | Flow | 10 µL/$10^6$ cell |
| Rabbit isotype control | Abcam | 172730 | Rabbit | Flow | 1:100 |
| Hoechst 33342 | SIGMA | B2261 | | IF | 1:12000 |

| | hPSC-vSMC + DMSO | hPSC-vSMC + MEKi | Adult AoSMC | Fetal AoSMC |
|---|---|---|---|---|
| **Average force (nN/post)** | 1.84 | 2.29 | 1.64 | 2.01 |
| **Standard deviation (nN)** | 0.50 | 0.53 | 0.51 | 0.60 |
| **Number of cells** | 76 | 75 | 130 | 40 |
| **Average Force Comparison with hPSC-vSMC + DMSO** | -- | Greater: $p < 0.0001$ | Lesser: $p < 0.005$ | Greater: not significant |
| **Average Force comparison with hPSC-vSMC + MEKi** | Lesser: $p < 0.0001$ | -- | Lesser: $p < 0.0001$ | Lesser: $p < 0.05$ |
| **Average Force comparison with Adult AoSMC** | Greater: $p < 0.005$ | Greater: $p < 0.0001$ | -- | Greater: $p < 0.001$ |
| **Average force comparison with Fetal AoSMC** | Lesser: not significant | Greater: $p < 0.05$ | Lesser: $p < 0.001$ | -- |

**Table S3**. Single cell force statistics. Summary statistics for the mPAD force data shown in Figure 5C. The categories hPSC-vSMC + DMSO and + MEKi pool all data from the corresponding treatments of JHU001-vSMC and WTC11-vSMC. The category Adult AoSMC combines data from lines AoSMC1, AoSMC2, and AoSMC3. The category Fetal AoSMC contains data only from line $AoSMC_f$. The significance of differences among cells from each category are represented by the p-value following a Welch's unpaired t-test comparison.

### *Other* Supplemental Information

**Supplementary Western Blot Figure Legends.**

(A) Western blots images showing tests for vSMC antibody specificity: Representative immunoblot exposures of entire protein transfers showing antibody staining for β-actin, ACTA2, CNN1, and TAGLN in cell lysates from JHU001. The COL1A1 antibody was not tested for specificity in this study. Each lane was loaded with 20 μg total protein. Lane 1: undifferentiated iPSCs; lane 2: iPSC-cardiomyocytes (CM1), differentiation day 35; lane 3: pooled vSMC samples treated with DMSO or with MEKi; lane 4: iPSC-CM2, differentiation day 35. JHU001-vSMC served as the positive control for smooth muscle markers, while undifferentiated JHU001-iPSCs and -CMs served as negative or low-expression controls.

(B) Western blot images for myosin heavy chain isoforms and vSMC markers in JHU001-vSMC following DMSO or MEKi treatment: All immunoblots (with representative exposures) of entire protein transfers showing expression of MYH9, MYH10, MYH11, ACTA2, CNN1, TAGLN, and β-actin in JHU001-derived vSMC treated with vehicle control DMSO or MEKi (as indicated). Molecular weight markers are shown on the left of each blot, and exposure times are indicated at the bottom. β-actin was used as the loading control. Red boxes indicate the bands selected for compositive presentation in the main figures (Fig. 1H and Fig. 2E).

(C) Schematic of SDS-PAGE and PVDF membrane showing membrane cutting strategy for protein analysis using Western blots: Samples prepared in Taiwan were loaded as three identical lane sets on a single SDS-PAGE gel, with each set separated by molecular weight markers (M). After electrophoresis, one membrane was stained with Coomassie Blue to visualize total protein loading for normalization. The replica sample sets were transferred to PVDF membranes for immunoblotting. Following transfer, the membranes were aligned using the molecular weight ladder and cut into defined molecular weight regions to allow probing with antibodies against target proteins of different sizes, including MYH11/COL1A1, ACTA2/CNN1, TAGLN (and SMTN, not shown). Dashed lines indicate the membrane cutting positions used to generate individual PVDF strips for separate antibody incubation.

(D) Whole membrane Western blot images of JHU001- and WTC11-derived vSMC lysates: Representative immunoblots showing COL1A1, CNN1, TAGLN, MYH11, and ACTA2 protein levels in JHU001- and WTC11-derived vSMC. Cells were cultured under SMCM or Lac medium (left) or were treated with Lac medium and cultured with DMSO or MEKi (right). Coomassie Blue staining is shown from parallel gels and used for normalization of membrane Set-1 and Set-2. Molecular weight marker masses are shown. Red boxes indicate the bands selected for composite presentation in the main figures, including Fig. 2E. Red arrowheads mark the expected molecular weight regions for the indicated target proteins. See (C) for membrane cutting scheme.

(E) Whole membrane Western blot images of H9-derived vSMC lysates: Representative images of immunoblots showing protein (COL1A1, ACTA2, CNN1and TAGLN) levels in H9-derived vSMC, treated with DMSO, Rap, or MEKi. Coomassie Blue staining of parallel gels is included as a total protein loading control. Molecular weight markers are indicated in kDa. Red boxes indicate the representative bands selected for presentation in the corresponding main figure panel, Fig. 6C. See (C) for membrane cutting scheme.

(F) Whole/cut membrane Western blot images of vSMC expression in AoSMCs following DMSO or MEKi treatment. Immunoblots showing β-tubulin, ACTA2, and TAGLN expression in AoSMC1 and AoSMC$_f$ cells treated with vehicle control DMSO or MEKi are provided. Note, the original membrane following protein transfer was cut (see dotted red line) prior to incubation with antibodies for β-tubulin and TAGLN. Other blots that were not cut are shown at the bottom of this image showing the individual protein signals. β-tubulin was used as the loading control. Molecular weight marker sizes are shown. Red boxes indicate the representative bands selected for presentation in the corresponding main figure panel, Fig. 4E.
